# Regulation of astrocyte lipid metabolism and ApoE secretion by the microglial oxysterol, 25-hydroxycholesterol

**DOI:** 10.1101/2022.05.03.490494

**Authors:** Anil G Cashikar, Danira Toral Rios, David Timm, Johnathan Romero, Justin M Long, Xianlin Han, David M. Holtzman, Steven M Paul

## Abstract

Neuroinflammation is a major hallmark of Alzheimer’s disease and several other neurological and psychiatric disorders and is often associated with dysregulated cholesterol metabolism. Relative to homeostatic microglia, activated microglia express higher levels of *Ch25h*, an enzyme that hydroxylates cholesterol to produce 25-hydroxycholesterol (25HC). 25HC is an oxysterol with interesting immune roles stemming from its ability to regulate cholesterol biosynthesis. Since astrocytes synthesize cholesterol in the brain and transport it to other cells via apolipoprotein E (ApoE)-containing lipoproteins, we hypothesized that secreted 25HC from microglia may influence lipid metabolism as well as extracellular ApoE derived from astrocytes. Here we show that astrocytes take up externally added 25HC and respond with altered lipid metabolism. 25HC increased extracellular levels of ApoE lipoprotein particles without altering *Apoe* mRNA expression, due to elevated *Abca1* expression via activation of LXRs and decreased ApoE reuptake due to suppressed *Ldlr* expression via inhibition of SREBP. Astrocytes metabolized 25HC to limit its effects on lipid metabolism via Cyp7b1, an enzyme responsible for 7α-hydroxylation of 25HC. Knockdown of *Cyp7b1* expression with siRNA prolonged the effects of 25HC on astrocyte lipid metabolism. 25HC also suppressed *Srebf2* expression to reduce cholesterol synthesis in astrocytes but did not affect fatty acid levels or the genes required for fatty acid synthesis. We further show that 25HC led to a doubling of the amount of cholesterol esters and their concomitant storage in lipid droplets. Our results suggest an important role for 25HC in regulating astrocyte lipid metabolism.

## INTRODUCTION

Neuroinflammation is an important hallmark of Alzheimer’s disease (AD) and other neurologic diseases as well as certain psychiatric disorders (1). Commonly recognized by the increased numbers of activated microglia and astrocytes in specific regions of the brain, neuroinflammation also entails dramatic changes in gene expression profiles of microglia and astrocytes. In AD, and in mouse models of AD, a specific activated state of microglia is commonly referred to as disease-associated microglia (DAM) (2–6). The gene expression signature of DAM shows down-regulation of homeostatic genes and up-regulation of several activation markers. One such activation marker is *Ch25h*, encoding cholesterol-25-hydroxylase. Single cell and single nuclei transcriptomics studies show myeloid cell specific overexpression of *Ch25h* in various mouse models of neurological disorders ((7); http://research-pub.gene.com/BrainMyeloidLandscape/BrainMyeloidLandscape2/#).

We have recently demonstrated that *Ch25h* is overexpressed in AD brain as well as in mouse models of amyloid deposition and tau-mediated neurodegeneration (8). Ch25h, the enzyme that hydroxylates cholesterol to 25-hydroxycholesterol (25HC) (9) is markedly upregulated by toll-like receptor (TLR) activation of macrophages (10, 11). Activation of primary microglia with the TLR-4 agonist, lipopolysaccharide (LPS) also upregulates *Ch25h* and results in increased synthesis and release of 25HC into the extracellular milieu (8). 25HC has been shown to have anti-viral properties and exhibit context-dependent immune-modulatory effects which can be either pro- or anti-inflammatory (12, 13). While loss of Ch25h and 25HC promoted IL1β production in mouse macrophages (14, 15), higher concentrations of 25HC amplified IL1β secretion in an ApoE-dependent manner in mouse microglia (8). More recently, we showed that 25HC disrupts hippocampal plasticity and learning in mice (16). These findings illustrate that 25HC not only exhibits autocrine effects on Ch25h-expressing myeloid cells but also supports paracrine effects on neighboring cells.

The paracrine effects of 25HC on astrocytes are not well understood, but most likely stem from its well-documented molecular actions, including binding to and activation of liver-X-receptors (LXRs) to promote lipid efflux (17), binding to INSIG-1 or INSIG-2 to suppress SREBP-mediated transcription of lipid synthesis genes (18, 19) and increasing cholesterol esterification through ACAT/SOAT enzymes (20, 21). Therefore, we posited that 25HC released from activated microglia may directly impact lipid metabolism in astrocytes and the secretion of lipoproteins containing ApoE.

Astrocytes are important for cholesterol synthesis in brain and are a major source of ApoE-containing lipoprotein particles central to cholesterol transport between astrocytes, neurons, and other cells (22). *APOE* is the most important risk factor for AD and contributes significantly to the risk of developing several other neurological and non-neurological disorders (23). A single copy of the *APOE-E4* allele confers a ∼ 3.7 times greater risk than the common variant, *APOE-E3*, whereas homozygous *APOE-E4/E4* carriers have a ∼12-15 times greater risk for developing AD. In a mouse model of amyloid deposition, ApoE has been reported to promote the localization of astrocytes to plaques (24). ApoE also markedly accelerates amyloid deposition (25, 26) as well as tau-mediated neurodegeneration in mouse models (27). Despite its pathophysiological roles in AD and other CNS diseases, the mechanisms underlying ApoE production and secretion in the brain are not clear. Here, we report that 25HC directly modulates astrocyte lipid metabolism via activation of LXR-mediated and inhibition of SREBP-mediated gene expression. As a result, 25HC promotes ApoE secretion via a post-translational mechanism. In addition, 25HC enhances esterification of cholesterol and the accumulation of lipid droplets.

## MATERIALS AND METHODS

### Chemicals

Oxysterols – 25HC (Cayman Chemical Company, 11097), 25-(C4 Top fluor^®^)-25HC (Avanti Polar Lipids, 810299P), 7α25diHC (Cayman Chemical Company, 11032), cholesterol (Avanti Polar Lipids, 700000), LPS (Sigma Aldrich, L2630), GSK2033 (Sigma Aldrich, SML1617), T0901317 (Cayman Chemical Company 71810). LY295427 was a kind gift from Dr. Douglas Covey at Washington University School of Medicine, St Louis, MO (61).

### Antibodies

Mouse anti-human ApoE monoclonal antibodies (HJ15.7 and biotinylated-HJ15.4; (62)) and mouse anti-mouse ApoE monoclonal antibodies (HJ6.2, HJ6.3 and biotinylated-HJ6.8; (63)) were from the Holtzman lab. Rabbit anti-Abca1 (Novus, NB400-105); Rabbit monoclonal anti-LDLR (EP1553Y) (Abcam, ab52818); Rabbit anti-ApoJ (Protein Tech, 12289-I-AP); Rabbit anti-Cyp7b1 (Invitrogen, PA5-75380); Mouse anti-β-Actin (Sigma, A5316); Goat anti-rabbit HRP secondary antibody (Leinco Technologies, R115); Goat Anti-rabbit Alexa 555 (Invitrogen, A32732); Goat Anti-mouse Alexa 488 (Invitrogen, A32723).

### Mice

Mice lacking *Ch25h* gene were originally developed in the lab of David Russell (10). Wildtype C57BL/6J (referred to as ‘wildtype’ or WT) and B6.129S6-*Ch25h*^tm1Rus^/J (referred to as ‘*ch25h-/-*’ or KO) mice were from Jackson Laboratories (Strain #016263). ApoE KI mice with the targeted replacement of the endogenous murine *Apoe* gene with human *APOE-ε3* or *APOE-ε4* were described earlier (31). All mice were maintained on C57BL/6J background and housed in Association for Assessment and Accreditation of Laboratory Animal Care (AAALAC) accredited facilities with *ad libitum* access to food and water on a 12-hour light/dark cycle. All animal procedures were approved by the Institutional Animal Care and Use Committee (IACUC) at Washington University and were in agreement with the AAALAC and WUSM guidelines.

### Cells

Primary astrocyte cultures were obtained from the cortex of neonatal mice (2-3 days old). Meninges were removed and cortices dissected in cold Hanks’ Balanced Salt solution without calcium and magnesium. The tissue was digested with 0.25% trypsin at 37°C for 5 min, and the cell suspension was filtered using a 100μm sterile filter. Cells were centrifugated at 300*g* for 5 minutes at 23 °C and resuspended in astrocyte medium (DMEM + 10% heat-inactivated fetal bovine serum + 1X Glutamax + 1X Sodium pyruvate and 1X Penicillin/Streptomycin – all from GIBCO). Cells were seeded in a T-75 flask previously coated with 10μg/ml poly-D-lysine (Sigma Aldrich, P7280) overnight at 37°C. Cells were cultured for 7-10 days with media changes every 3-4 days to reach a confluent layer of astrocytes. For astrocyte cultures, loosely attached cells microglia were shaken off, followed by trypsinization and replating into desired tissue culture plates at a density of 2 X 10^5^ cells/ml. All treatments were conducted on astrocyte serum-free media (DMEM/Neurobasal (1:1) + 1X Glutamax + 1X Sodium pyruvate + 0.1% of fatty acid-free bovine serum albumin and 1X penicillin/streptomycin – all were from GIBCO). For 25HC treatment, an identical volume of ethanol was added as Vehicle. For microglia cultures, the procedure was similar except the culture media also contained 5ng/ml GM-CSF (GoldBio # 1320-03-5) to stimulate microglial proliferation. When large numbers of microglia were observed floating, the flasks were shaken at 200rpm for 30 minutes at 37°C. The floating cells were collected by centrifugation and the cells were resuspended in microglia media (DMEM/F12 + 1X Glutamax + 1X Sodium pyruvate + 0.1% of fatty acid-free bovine serum albumin and 1X penicillin/streptomycin + 1X ITS supplement (R&D Systems AR013) + 25 ng/mL M-CSF (GoldBio 1320-09-10). All treatments were done in the same serum-free medium.

### Fluorescent 25HC uptake

Cells were seeded at a density of 4 X 10^4^ cells/ well in 8-Well chambered #1.5 coverglass (CellVis, C8-1.5P). A stock of 2mM 25-(C4 Top fluor^®^)-25HC was prepared in ethanol. Astrocyte treatment media containing 1.8μM unlabeled 25HC (90%) and 0.2μM topfluor-25HC (10%) was prepared to result in a total 25HC concentration of 2μM. Astrocyte culture media was replaced with treatment media containing 25HC. A kinetic study of 25HC uptake was conducted by fixing cells at 30 min, 1h, 2h, 4h, 1 day with paraformaldehyde (Electron microscopy Sciences, Cat. No. 15710). Cells were counterstained with DAPI and imaged on Zeiss LSM880 using identical settings at the Washington University Center for Cellular Imaging. Images were processed similarly using ImageJ (Fiji).

### 25HC quantitation

The amount of 25HC released into conditioned culture media by astrocytes and microglia was quantified using previously described methods (29). Briefly, conditioned media was subjected to derivatization with dimethyl glycine followed by liquid chromatography-mass spectrometry. The amount of 25HC was estimated based on an internal standard of deuterated-25HC.

### Drug treatments

Mouse primary astrocytes were plated at a density of 2 X 10^5^ cells/ml in a poly-D-lysine coated multi-well plate as needed. Cells were allowed to attach for 2 days, media were removed, and the treatments were diluted in astrocyte serum-free media. Stock solutions of 25HC, cholesterol, T0901317, and LY295427 were made in ethanol and for GSK2033 in dimethyl sulfoxide (DMSO) at 10mM final concentration. 25HC treatments were at 2μM final concentration (5000-fold dilution of the stock solution for all experiments. For vehicle control samples, an identical volume of ethanol was added resulting in a final ethanol concentration of 0.02% (∼3mM). For the 25HC concentration series, the highest concentration tested was 4μM and corresponding amounts of ethanol as vehicle was used as control. In experiments to test how 25HC influences LXR- and SREBP-pathways, LY295427 and GSK2033 were added to astrocyte serum-free media at a final concentration of 5μM. For GSK2033, DMSO was the vehicle control. Treatments were terminated after 1, 2, or 4 days by collecting media and cells for downstream processing as needed.

### RNAi

Short interfering RNA (siRNA) for Silencer Select siRNA for mouse *Cyp7b1* gene (siRNA ID: s64780; Cat. No. 4390771) and a Silencer Select Negative control siRNA (Cat. No.4390843) were obtained from Ambion (ThermoFisher Scientific). Astrocytes were plated at a density of 2 X 10^5^ cells/well in a 12-well plate. RNAi was transfected into primary astrocytes at 80-90% confluence using Lipofectamine RNAiMax (Invitrogen by Thermo Fisher, Cat. No. 56531). Knockdown efficiency was confirmed with qPCR as well as western blotting. Treatment of siRNA transfected astrocytes with drugs was done 24h after siRNA transfection.

### ApoE ELISA

ELISA for mouse and human ApoE was performed as described earlier (31, 63). The capture antibody was HJ6.2 and HJ15.7 the detection antibody was biotinylated-HJ6.8 and biotinylated HJ15.4 for quantifying mouse and human ApoE, respectively. The ELISA plates were coated with 10μg/ml HJ15.7 (human ApoE) or 5μg/ml HJ6.2 (mouse ApoE) for 2h at room temperature. The plates were washed with PBS and blocked with 2% BSA in PBS. Samples were diluted appropriately in 1% BSA in PBS with protease inhibitors, loaded into the plates in duplicates and incubated overnight at 4°C. The next day, the plates were washed with PBS, followed by addition of the appropriate detection antibody at 50ng/ml - biotinylated-HJ6.8 (mouse ApoE) or biotinylated HJ15.4 (human ApoE) and incubated for 1h at room temperature. The plates were washed and incubated with streptavidin-poly-HRP40 (Fitzgerald #65R-S104PHRP) for 1h at room temperature. The plates were then developed with Super Slow TMB (Sigma-Aldrich, Cat# T5569). The development was stopped using 2N sulfuric acid and the plate was read at 450nm.

### Immunoblotting

For secreted experiments, media were collected after the treatment. Cells were washed with PBS and lysed with RIPA buffer (Millipore, Cat. No. 20-188) containing ProBlock Gold™ Mammalian protease inhibitor cocktail (Goldbio, GB-331-1) and Benzonase (Sigma, E1014-5KU). Samples were run on 4-15% Mini protean TGX stain free gels (Biorad, Cat. No.4568085) at 100V for 1h. Gels were transferred to PVDF membranes at 100V by 1 h. Membranes were blocked with a 5% non-fat dry milk in tris-buffered saline with 0.1% Tween-20 (TBST). Primary antibody was diluted in blocking buffer and incubated at 4°C overnight. Blots were washed and incubated with the secondary antibody conjugated to HRP (Leinco Technologies, M114). Blots were developed with chemiluminescent development using Immobilon ECL Ultra Western HRP substrate (Millipore, CS222617) and imaged using the ChemiDoc System (Bio-Rad Labs). Band intensities were quantified using the Image Lab software (Bio-Rad Labs).

### ApoE non-denaturing gradient gel electrophoresis

Cells were plated in a 6-well plate with a total density of 4 X 10^5^ cells/well. Supernatants were collected 4 days after treatment. Protein was concentrated from the media using VivaSpin Turbo 4 (10000 MWCO) concentrator tubes (Sartorius, VS04T01). Protein samples were run on 4-15% Mini protean TGX stain-free gels (Bio-Rad) at 100V for 4h at 4°C. Gels were transferred to PVDF membrane at 100V for 1h at 4°C. Immunoblotting was carried out as described above.

### qPCR

RNA was isolated using the Quick-RNA Miniprep Plus Kit (Zymo Research, Cat. No. R1058). cDNA was synthesized from total RNA using High-Capacity cDNA Reverse Transcription kit with RNase inhibitor (Applied Biosystems, Cat No.4374966 Foster City, CA, USA). For the qPCR reaction mix, PrimeTime probe-based qPCR assays (see Table I) and PrimeTime Gene Expression Mastermix (Cat. No.1055771) were obtained from IDT (Integrated DNA Technologies, Inc). qPCR reactions were run using the Fast mode on a QuantStudio™ 3 Real-Time PCR Instrument (Applied Biosystems by ThermoFisher, A28131). For gene expression experiments, data were normalized against actin (Actb) and the 2^−ΔΔCt^ method was used to calculate relative gene expression value (Relative Quantity, RQ). For absolute quantification of *Ch25h* expression in microglia and astrocytes, PrimeTime probe-based qPCR assay for *Ch25h* (Cat. No. Mm.PT.58.42792394.g) from IDT was used to determine a standard curve for threshold cycle (C_T_) values against the number of copies of *Ch25h* cDNA. Since *Ch25h* is an intronless gene, the plasmid construct with the mouse *Ch25h* gene in pcDNA3.1 (Genscript, OMu18523D) was used to define the standard curve. The C^T^ values obtained for cDNA prepared from 200ng RNA isolated from equal number (2 X 10^5^) of microglia or astrocytes were interpolated against the standard curve to determine the absolute number of copies of *Ch25h* in microglia and astrocytes.

**TABLE I.**
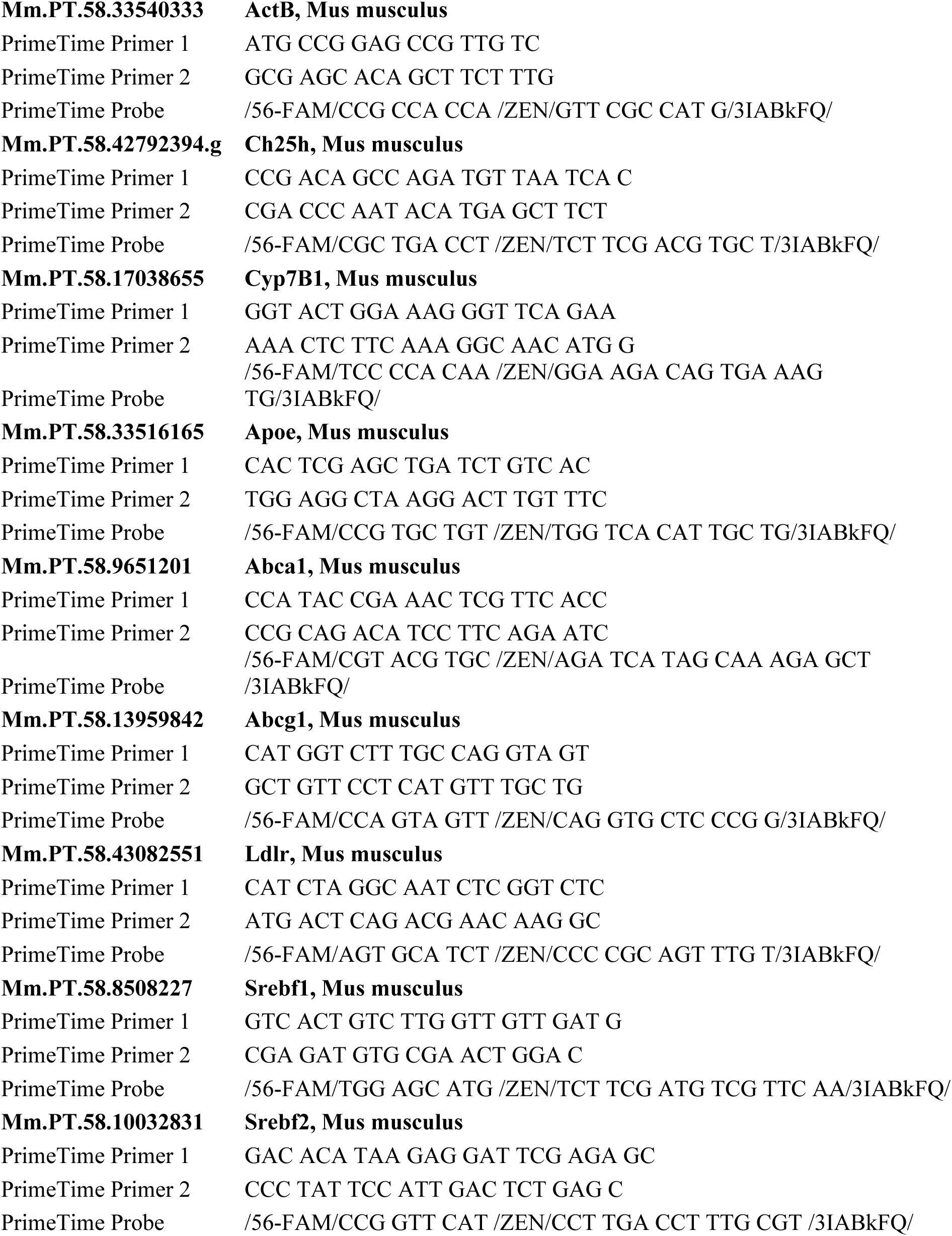
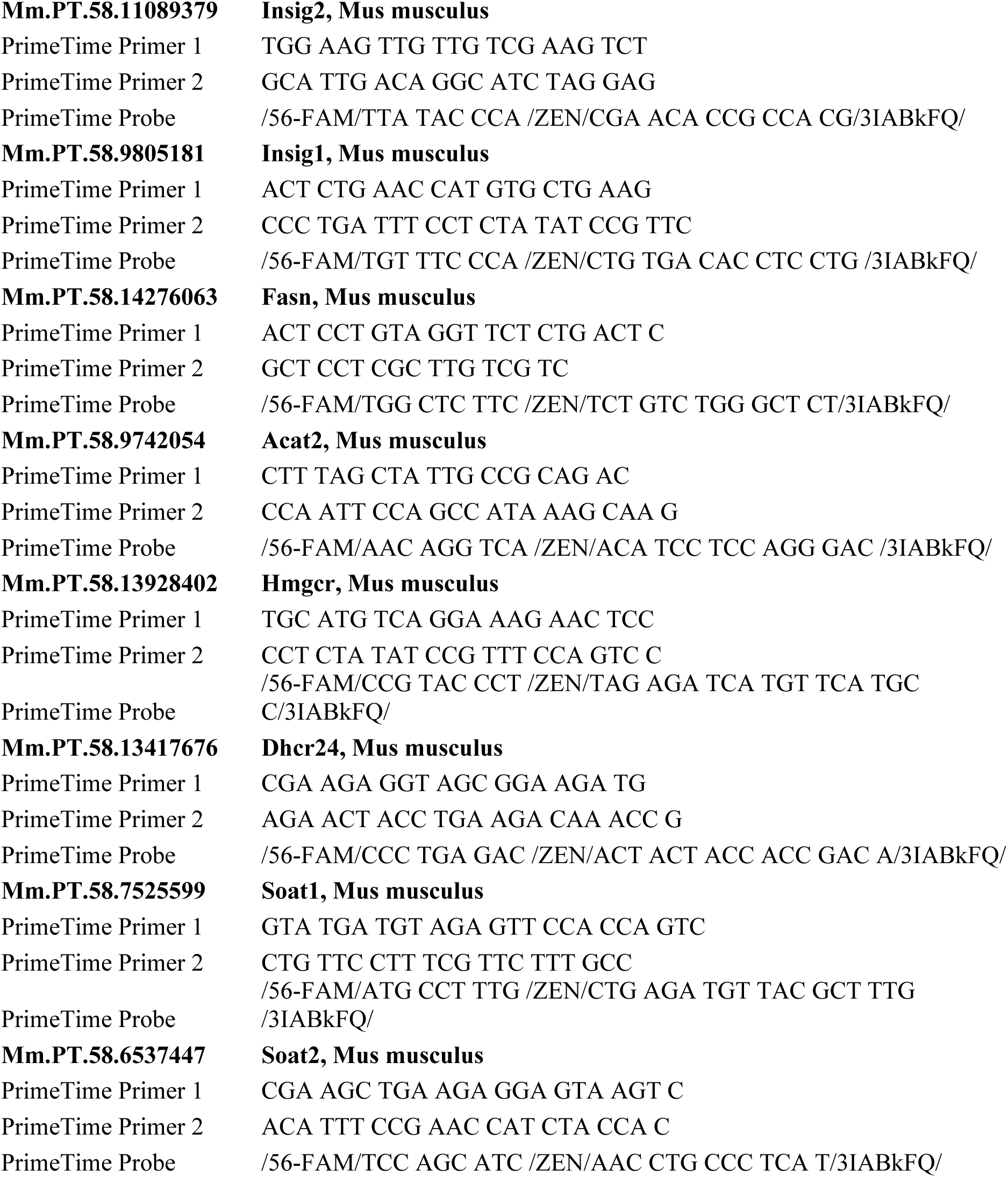
Probe-based qPCR assays from IDT (Integrated DNA Technologies)

### Lipid droplet analysis

Mouse primary astrocytes were seeded at a density of 3 X 10^4^ cells in a 8-well chamber slide (CellVis, C8-1.5P). Oleic acid was added as Oleic acid-bovine serum albumin (Sigma O-3008) and used with cell culture BSA (Sigma A-1595) as vehicle control. To address the ability of 25HC to induce lipid droplets, the following treatments were added separately or in combination in astrocyte serum-free media: Vehicle (ethanol + BSA), 25HC (2μM), oleic acid (30μM), and the acyl-coenzyme A: cholesterol acyltransferase (SOAT) inhibitor, avasimibe (1 μM; APExBIO cat#A4318). After 24 hours, the cells were washed, fixed in 4% paraformaldehyde and stained with BODIPY 493/503 (Thermo Fisher Scientific, D3922) and DAPI. Images from three independent experiments were acquired on the Nikon Spinning Disk Confocal microscope at the Washington University Center for Cellular Imaging. Lipid droplets and cells counts were analyzed using ImageJ (Fiji).

### Lipidomic analysis

1 X 10^6^ astrocytes were grown in six 6-cm dishes and treated with Vehicle or 25HC (2μM). After 2 days, the media were collected and centrifugated at 4000 rpm x 5 min to remove cell debris. The cells were washed and collected in DPBS 1X. Cells were transferred into a microtube and the supernatant was removed after centrifugation at 10000 g X 5 min at 4°C. Samples were frozen on dry ice and stored at -80C. Lipidomic analysis was carried out at the Functional Lipidomics Core at Barshop Institute for Longevity and Aging Studies of the University of Texas Health Science Center, San Antonio, TX. Briefly, lipid extracts of cultured cells were prepared by the procedure of Bligh-Dyer extraction (64) and a premixed solution of internal standards was added based on the protein content of cell homogenates. Individual molecular species of free fatty acids were identified and quantified after derivatization as previously described (65). Total cholesterol and free cholesterol were quantified after derivatization with methoxyacetic acid as described (66). The mass levels of individual cholesterol ester species were calculated from total cholesteryl ester mass and their composition was profiled using precursor-ion scanning of m/z 369. Data processing based on the principles of shotgun lipidomics such as selective ionization, low concentration of lipid solution, and correction for differential isotopologue patterns and kinetics of fragmentation was conducted as previously described (67).

### Statistical methods

To determine statistical significance (p value of <0.05 was considered significant; * 0.05-0.01; **0.01-0.001; ***<0.001). Student’s t test or two-way ANOVA with appropriate post hoc multiple comparisons tests (as indicated) were carried out with the help of GraphPad Prism version 9.0.

## RESULTS

### Inflammatory stimuli induce secretion of 25HC in microglia, but not astrocytes

Multiple studies reporting single cell-RNAseq data have shown that Ch25h is predominantly (perhaps exclusively) expressed in microglia in the brain. However, expression of Ch25h has also been reported in primary cultured astrocytes (28) and may indicate an altered astrocyte gene expression profile upon isolation and culture. To validate microglia-specific expression of Ch25h, we first tested for purity of primary mouse microglia and astrocytes using immunofluorescence with antibodies specific to Iba1 and GFAP, respectively (Sup Fig 1A & B). While all cells in the microglia culture were positive for Iba1, about 1% of cells in astrocyte cultures were Iba1-positive microglia. Next, we measured the expression of Ch25h in cultured primary mouse microglia and astrocytes using a qPCR method for absolute quantitation. A standard curve was constructed with cycle threshold (C_T_) value against gene copy number using a known concentration of purified plasmid DNA (pCDNA3.1 with mouse *Ch25h* gene) as shown in Sup Fig 1C. Microglia and astrocytes were challenged with 100ng/ml LPS to stimulate transcription of inflammatory genes including Ch25h. Equal amounts of RNA (120ng) from microglia and astrocytes were used to prepare cDNA for qPCR. The C_T_ values for the samples were compared against the standard curve to determine the expression of Ch25h between the two cell types. In microglia, Ch25h was low in the absence of LPS and increased about 100-200-fold after treatment with LPS (Fig 1A). In astrocytes, Ch25h was low in the absence of LPS and increased only marginally upon treatment with LPS, presumably resulting from the low levels of contaminating microglia (Fig 1A).

**Figure 1:**
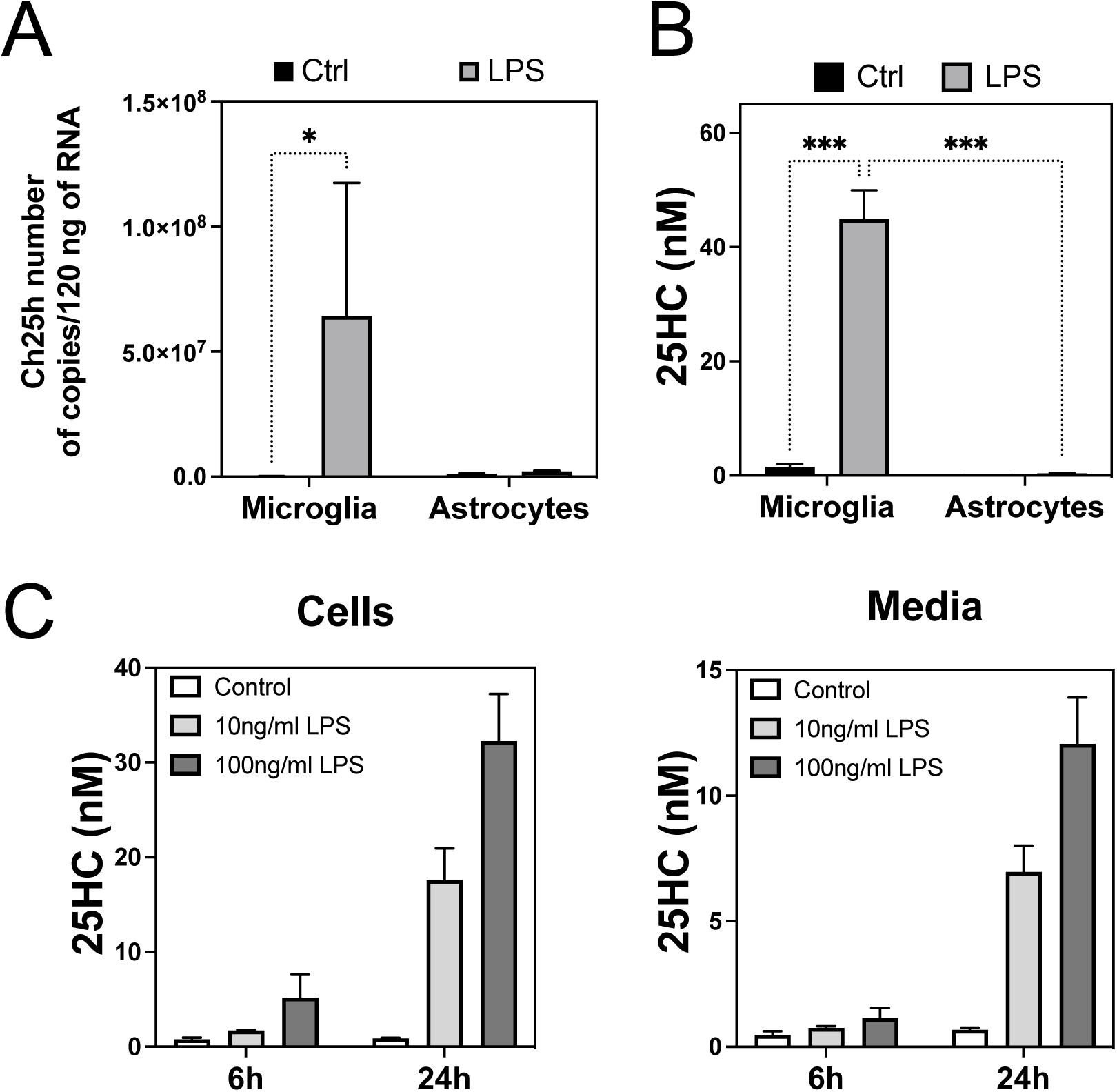
Microglia express Ch25h and secrete 25HC in response to LPS treatment. (A) Comparison of Ch25h expression between mouse microglia and astrocytes by qPCR in response treatment with 100ng/ml LPS for 24h. Data is shown absolute copy numbers of Ch25h mRNA per 200,000 cells. A plot of CT values versus copy number is provided in Supplementary Figure 1. (p-values, * < 0.05; two-way ANOVA) (B) Measurement of 25HC in conditioned culture media from microglia or astrocytes with or without treatment with 100ng/ml LPS, as indicated. (p-values, *** <0.0002, Two-way ANOVA with Tukey test for multiple comparisons). (C) Comparison of amount of 25HC in cells versus media at 6 or 24h after treatment of wildtype microglia with vehicle (control), 10ng/ml LPS or 100ng/ml LPS. For direct comparison of the distribution of 25HC, cell extracts were prepared in a volume proportional to the volume of media used for cell culture (200,000 cells in 1ml media).

We next examined the synthesis of 25HC by microglia and astrocytes with or without LPS. Cells were treated with LPS for 24h as before. Conditioned culture media was collected, debris precleared by centrifugation and the media supernatants were subjected to derivatization by dimethyl glycine followed by liquid chromatography and mass spectrometry-based quantitation as described previously (29). While microglia produced high levels of 25HC after LPS treatment, astrocytes produced no appreciable amounts of 25HC with or without LPS (Fig 1B). High levels of 25HC produced by LPS-treated microglia reflected the high levels of Ch25h expression, demonstrating a causal relationship between Ch25h expression and 25HC production in microglia. Therefore, we conclude that inflammatory stimuli triggered overexpression of Ch25h and production of 25HC by microglia, but not astrocytes. A very low level of Ch25h expression could be observed in our primary cultures of isolated neonatal mouse astrocytes most likely due to contaminating microglia. However, no 25HC production and secretion was detected from astrocytes with or without endotoxin stimulus.

It has been suggested that 25HC may be synthesized by enzymes other than Ch25h (30). To test this, we examined 25HC synthesis after LPS challenge in microglia from wild-type and *ch25h-/-* mice (Sup Fig 1D). Increasing amounts of 25HC as a function of time were observed exclusively in media from wild-type microglia, whereas no 25HC was detectable in media from *ch25h-/-* microglia. Thus, microglia from *ch25h-/-* mice produce no 25HC as previously demonstrated (8).

We next examined the proportion of 25HC that was secreted into the culture media relative to that retained within microglial cells. To do this, we prepared cell extracts after collecting the conditioned media. The volume of extraction buffer used was equal to the volume of the conditioned culture media to facilitate a direct comparison of 25HC levels. About 30-40% of 25HC produced by microglia after exposure to LPS was released extracellularly into the media while retaining 60-70% of it within cells (Fig 1C). This distribution is in agreement with that observed earlier for ApoE2 and ApoE4-expressing microglia (8). This suggests that microglial 25HC not only elicit autocrine effects within microglia but may also exert paracrine effects on surrounding cells such as neurons, astrocytes, and other cells to facilitate inter-cellular crosstalk.

### Astrocytes internalize 25HC

To test the hypothesis that the 25HC secreted by microglia may influence other cells, we first tested whether cultured astrocytes could take up and internalize 25HC. Astrocytes were treated with 2μM 25HC mixed with 10% (0.2μM) BODIPY-labeled fluorescent 25HC (TopFluor-25HC) in culture media, and the uptake of fluorescent 25HC was measured over a period of 24 hours (Fig 2A). Very little fluorescence, if any, could be detected at 2h. By 4h, a general increase in fluorescence intensity outlining the cells was visible suggesting that the 25HC was present in the plasma membrane. By 24h, 25HC was clearly present in intracellular vesicles in virtually every cell. To confirm whether astrocytes were able to take up unlabeled 25HC added externally into the culture media in the same timeframe, we quantified 25HC remaining in the culture media after 24 hours of contact with astrocytes. Interestingly, when astrocytes were treated with 1μM or 2μM 25HC, only 44.9nM (± 1.1nM) and 84.0nM (± 8.6nM) 25HC remained in the conditioned media after 24 hours (Fig 2B). These results suggest that nearly 95% of the externally added 25HC was readily taken up by astrocytes within 24h.

**Figure 2:**
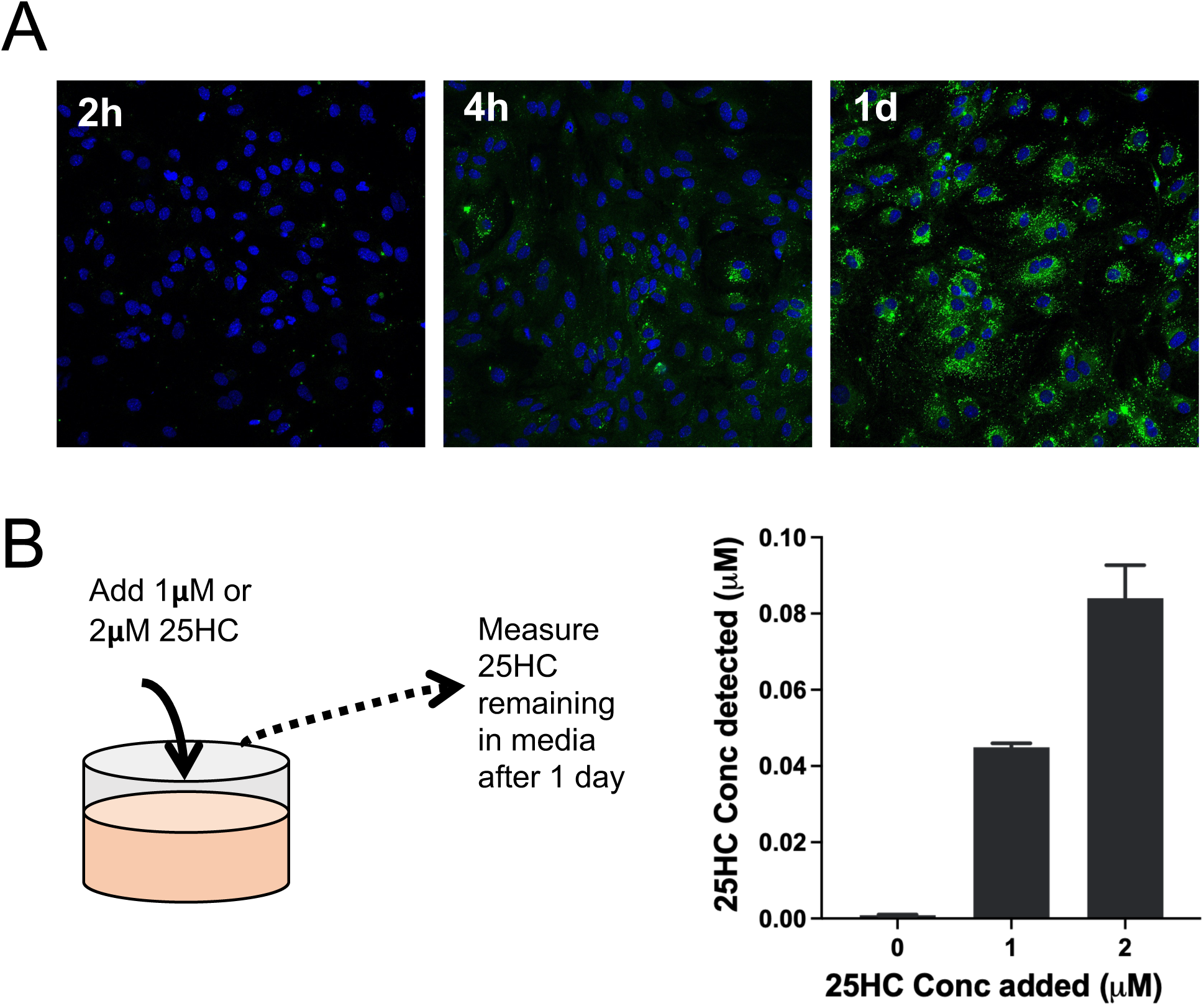
Uptake of 25HC by astrocytes. (A) Fluorescence microscopy of astrocytes treated with 2μM 25HC (containing 25HC (1.8μM) and TopFluor-25HC (0.2μM)) for various times. Time-dependent increase in fluorescence intensity of cells within intracellular vesicles was observed within 1 day. (B) On the left is a schematic of the experiment showing the addition of 25HC to a final concentration of 1μM or 2μM to astrocytes. Uptake of 25HC by astrocytes was monitored using LCMS to measure 25HC remaining (detected) in the culture media after 1 day of incubation on the right.

### 25HC stimulates ApoE secretion from astrocytes

Since ApoE is a major lipid carrier in the brain produced by astrocytes and its levels and secretion are thought to be central to AD pathogenesis, we tested whether 25HC impacts ApoE secretion from astrocytes. Astrocytes were incubated with 25HC, cholesterol, or vehicle (ethanol) and the ApoE in the culture media accumulating over 48 hours was measured by ELISA (Fig 3A). While cholesterol or ethanol (vehicle control) had no effect on ApoE secretion by astrocytes, 25HC increased ApoE secretion by about 200%. The stimulatory effect of 25HC was comparable to an established LXR agonist such as T0901317.

**Figure 3:**
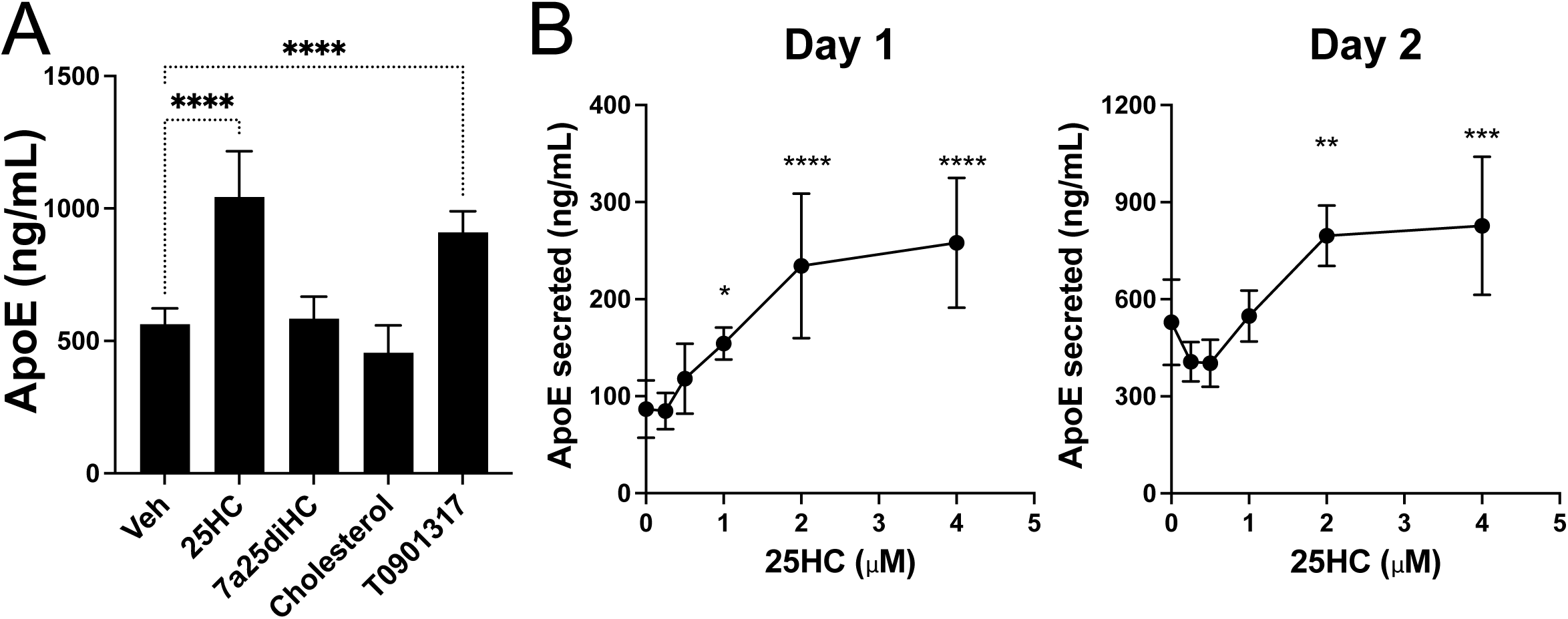
25HC increases extracellular ApoE in astrocytes. (A) Astrocytes were incubated with 2μM final concentration of 25HC, 7α25diHC, cholesterol, T0901317 or with an appropriate amount of vehicle (ethanol). ApoE was measured by ELISA in the conditioned media after 2 days. (p-values *<0.0332; ****<0.0001 – Two-way ANOVA with Sidak’s multiple comparisons test) (B) Effect of 25HC concentration and time on the accumulation of extracellular ApoE from astrocytes. (p-values * 0.05-0.01; **0.01-0.001; ***<0.001; ****<0.0001 – Two-way ANOVA with Dunnett’s multiple comparisons test).

We next tested the effect of 25HC at different concentrations over time. 25HC increased ApoE secretion from astrocytes in a concentration- and time-dependent manner. At earlier time points (1-day), basal secreted levels of ApoE in conditioned media were low under control conditions and increased by 4-5 fold over 2 days (Fig 3B). The amount of extracellular ApoE increased 2-3-fold following treatment with 25HC (2μM) relative to basal levels at both time points.

### 25HC does not stimulate expression of ApoE mRNA

To better understand the cellular mechanisms responsible for the increased extracellular ApoE in 25HC-treated astrocytes, we isolated RNA from astrocytes treated for 1 day and examined *Apoe* mRNA levels by qPCR. 25HC failed to alter the expression of *Apoe* mRNA at any time. Treatment of astrocytes with vehicle (ethanol) or cholesterol did not alter the expression of *Apoe* mRNA (Fig 4A). A synthetic non-steroidal LXR agonist, T0901317, increased *Apoe* expression by 1.5-fold. Longer treatment with 25HC for 4 days also did not alter *Apoe* gene expression (Fig 4B).

**Figure 4:**
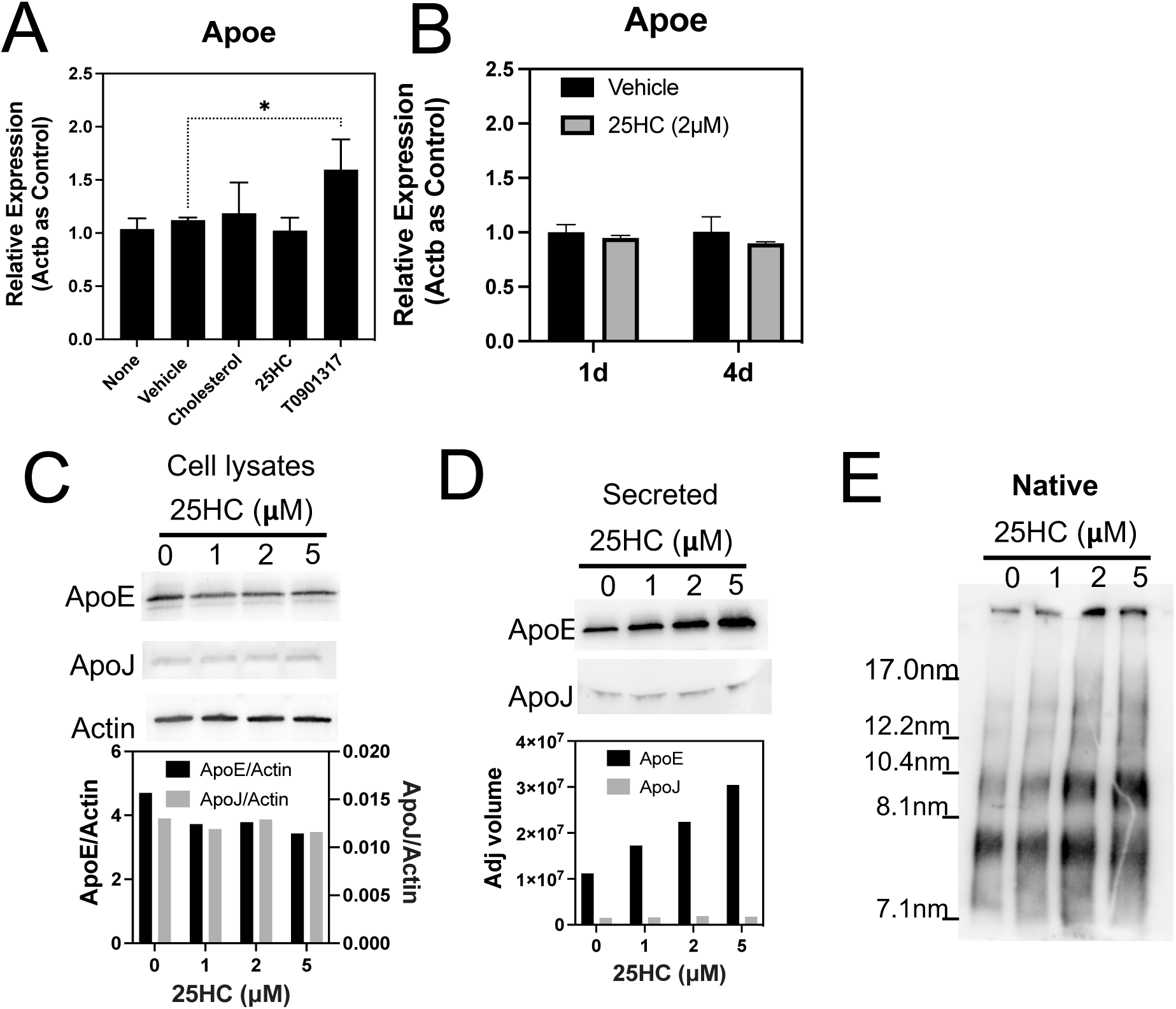
25HC increases extracellular ApoE without altering *Apoe* mRNA expression. (A) Expression of *Apoe* mRNA was measured by qPCR in astrocytes treated with 2μM each of cholesterol, 25HC or T0901317 or an appropriate amount of vehicle (ethanol) or without any treatment (none). Data is normalized to an *Actb* (actin B) as the endogenous control gene and shown as relative expression compared to ‘None’. (*0.05-0.01; Two-way ANOVA with Dunnett’s multiple comparisons test). (B) Expression of *Apoe* mRNA was measured by qPCR in astrocytes treated with 2μM 25HC (grey bars) or vehicle (black bars) for 1 day (1d) or 4 days (4d). (C) Immunoblots of cell lysates from astrocytes treated with 0, 1, 2 or 5μM 25HC for 48 hours. Antibodies for ApoE, ApoJ or actin are shown on the left of corresponding blots. Bar graphs below the blots show band quantitation after baseline correction normalized for actin band intensity. (D) Immunoblots of conditioned media from astrocytes treated with 0, 1, 2 or 5μM 25HC for 48 hours. Antibodies for ApoE or ApoJ are shown on the left of corresponding blots. Bar graphs below the blots show band quantitation after baseline correction. (E) Immunoblot for ApoE of conditioned media from astrocytes treated with 0, 1, 2 or 5μM 25HC for 48 hours, and samples were run on native polyacrylamide gel electrophoresis.

We further examined the amount of ApoE in cell lysates by immunoblotting and compared this with a control protein such as actin B or another apolipoprotein, ApoJ. None of these other astrocytic proteins were altered by 25HC when measured in cell lysates (Fig 4C). We next tested conditioned media for secreted ApoE and ApoJ. 25HC increased secretion of ApoE but not ApoJ suggesting that the effect was specific for ApoE (Fig 4D). Further, we did not observe any difference in ApoE expression or secretion between wildtype or ch25h-/-astrocytes (data not shown). These observations show that the observed increase in the levels of ApoE secreted into media following 25HC treatment occurs without substantially altering intracellular levels of ApoE.

Next, we examined whether the increased secretion of ApoE by 25HC was associated with an alteration in ApoE lipidation. To this end, we concentrated the variously treated conditioned media and separated the secreted ApoE by non-denaturing (native) gel electrophoresis followed by immunoblotting as described previously (31). While each band on the native blot intensified with increasing 25HC concentration, the relative proportions of each of the bands were not different between 25HC-treated and vehicle-treated samples (Fig 4E). This suggests that the extracellular ApoE secreted following 25HC treatment of astrocytes was not associated with a change in lipidation.

### 25HC upregulates Abca1 and downregulates Ldlr expression

We next assessed the key genes responsible for ApoE efflux and reuptake by astrocytes. As a relatively weak ligand of the liver-X-receptor (LXR), 25HC may increase the expression of *Abca1*, a gene controlled by LXRs. 25HC is also a strong suppressor of the sterol regulatory element binding protein (SREBP)-mediated transcription of genes including *Ldlr*. Abca1 is central to the efflux of ApoE lipoproteins while Ldlr facilitates its reuptake into cells (32). By qPCR, we observed that 25HC increased *Abca1* expression after 1 day of treatment by about 2.5-fold (p< 0.01), similar to a 3-fold increase observed with the synthetic LXR agonist, T0901317 (Fig 5A). Vehicle (ethanol) had no significant effect on *Abca1* expression. On the other hand, 25HC strongly suppressed the expression of *Ldlr* to about 10% of control (p < 0.001), while treatments with vehicle or T0901317 showed no effect on *Ldlr* expression (Fig 5B). We confirmed the effects of 25HC on Abca1 and Ldlr mRNA expression by immunoblotting for the respective proteins. Levels of Abca1 protein were increased following 25HC treatment, whereas Ldlr protein levels were decreased following 25HC treatment (Fig 5C). Levels of actin B and ApoE were unaffected in cell lysates by 25HC treatment.

**Figure 5:**
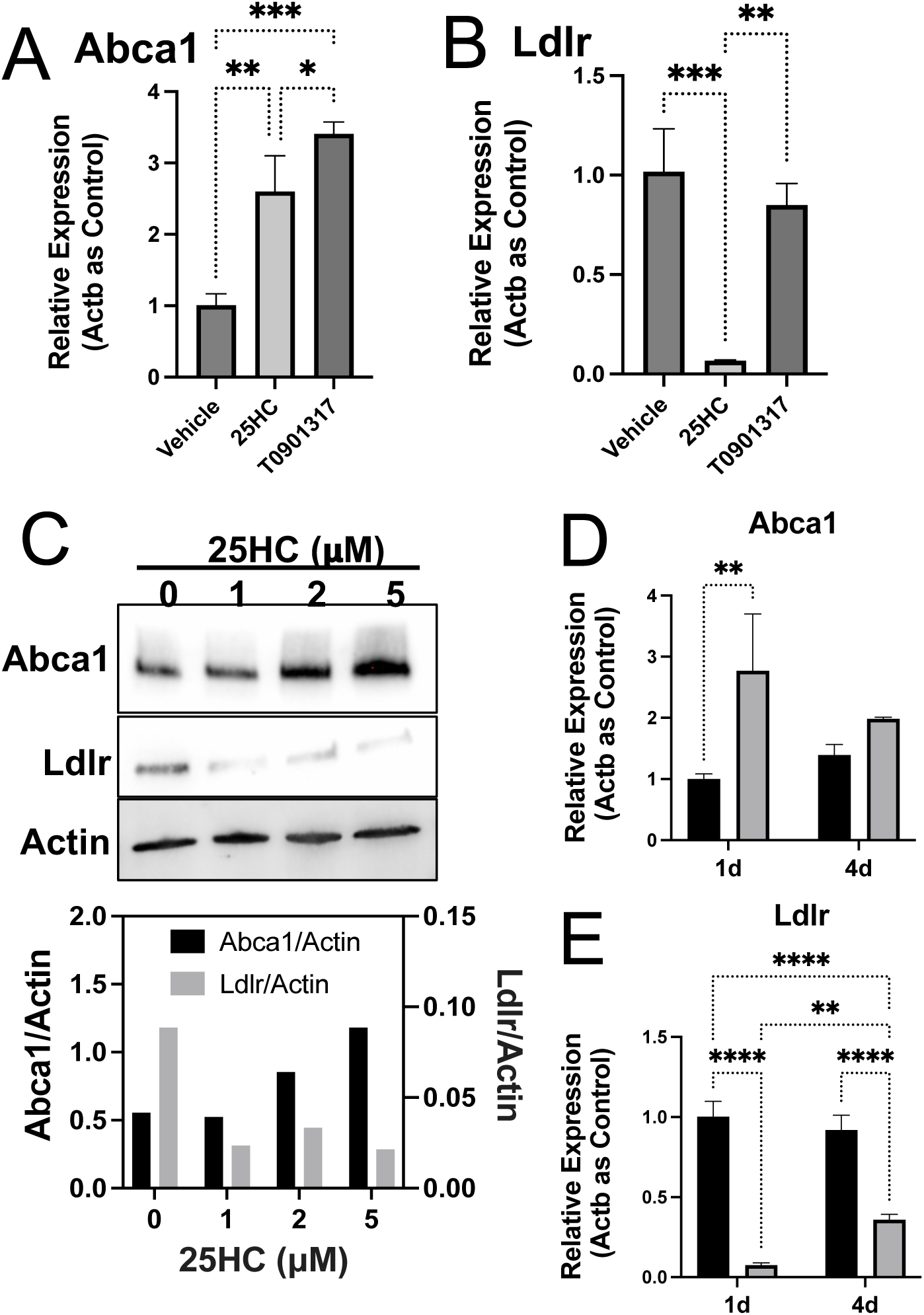
25HC modulates Abca1 and Ldlr mRNA and protein levels in astrocytes. (A & B) Expression of *Abca1* (A) and *Ldlr* (B) mRNA in response to treatment of astrocytes with vehicle, 25HC (2μM) or T0901317 (2μM) was assessed by qPCR. Data is normalized to *Actb* (actin B) as the endogenous control gene and shown as relative expression compared to ‘vehicle’. (*0.05-0.01; **0.01-0.001; ***<0.001; ****<0.0001; Two-way ANOVA with Sidak’s multiple comparisons test). (C) Immunoblots of cell lysates from astrocytes treated with 0, 1, 2 or 5μM 25HC for 48 hours. Antibodies for Abca1, Ldlr or actin are shown on the left of corresponding blots. Bar graphs below the blots show band quantitation after baseline correction and normalized for actin band intensity. (D & E) Effect of 25HC treatment of astrocytes over 4 days (grey bars) relative to 1 day (black bars) on the expression of Abca1 (D) and Ldlr (E) measured by qPCR as described in A and B. (*0.05-0.01; **0.01-0.001; ***<0.001; ****<0.0001; Two-way ANOVA with Sidak’s multiple comparisons test)

To test how long 25HC’s effect lasts on the expression of *Abca1* and *Ldlr* in astrocytes, we compared astrocytes treated with 25HC for 1 day and 4 days. While 25HC increased *Abca1* expression by about 2.5-fold after 1 day, we observed only an insignificant change on day 4 (Fig 5D). Likewise, 25HC reduced *Ldlr* expression to about 10% on day 1, but by day 4 *Ldlr* expression was approximately 35% of control (p<0.0001) (Fig 5E).

### Astrocytic Cyp7b1 metabolizes 25HC to reduce 25HC effects on lipid metabolism

To better understand the factors regulating 25HC’s effect on gene expression in astrocytes, we examined whether 25HC was being actively metabolized. 25HC is metabolized to 7α, 25-dihydroxycholesterol (7α25diHC) by Cyp7b1, a cytochrome P450 enzyme (Fig 6A). Unlike 25HC, 7α25diHC has no effect on ApoE secretion from astrocytes (Fig 3A). Interestingly, brain cell RNAseq data shows that while *Ch25h* is primarily expressed by microglia, *Cyp7b1* is primarily expressed in astrocytes and oligodendrocytes (33, 34). We hypothesized that if 25HC taken up by astrocytes is further metabolized by Cyp7b1 then the latter may serve to limit the effects of 25HC. To test this, we knocked down *Cyp7b1* expression in astrocytes using short interfering RNA (siRNA). We first ensured that siRNA against *Cyp7b1* was effective using qPCR (Fig 6B) and western blotting (Fig 6C). In *Cyp7b1* siRNA transfected astrocytes, *Cyp7b1* expression was reduced by about 90%. There was no significant effect of 25HC on *Cyp7b1* expression in astrocytes transfected with control or *Cyp7b1* siRNA (Fig 6B). Immunoblotting showed that Cyp7b1 protein levels were also markedly reduced in astrocytes transfected with *Cyp7b1* siRNA relative to a control siRNA (Fig 6C).

**Figure 6:**
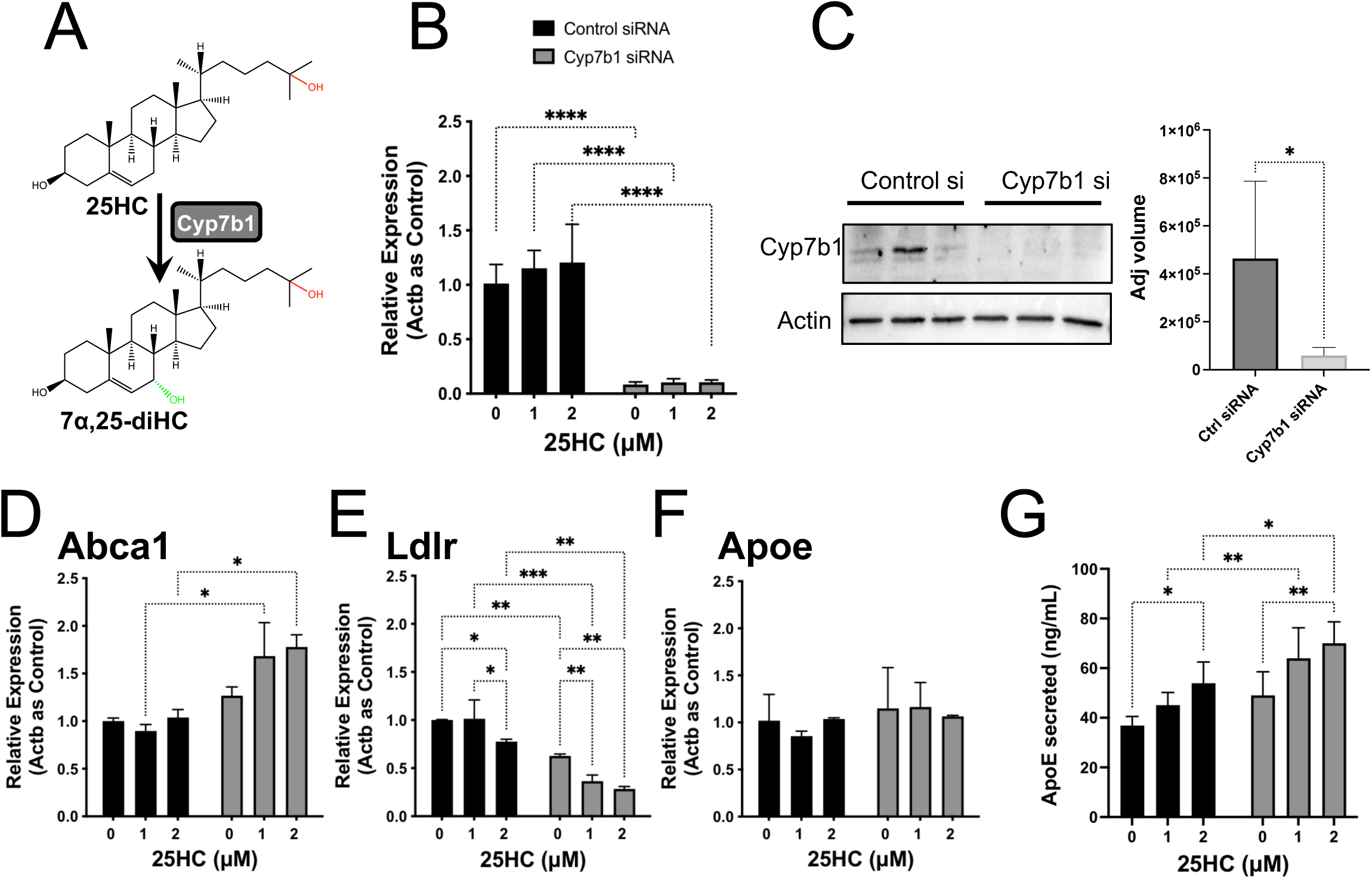
Astrocyte-expressed Cyp7b1 limits the effect of 25HC on extracellular ApoE levels and lipid metabolism in astrocytes. (A) Hydroxylation of 25HC to 7α,25-diHC by the 7α-hydroxylase enzyme, Cyp7b1. (B) Expression of *Cyp7b1* gene assessed by qPCR in astrocytes transfected with control siRNA (black bars) or Cyp7b1 siRNA (grey bars) followed by treatment with 0, 1μM or 2μM 25HC for 2 days. (****<0.0001; Two-way ANOVA with Sidak’s multiple comparisons test). (C) Immunoblots for Cyp7b1 and actin in astrocytes transfected with control siRNA or Cyp7b1 siRNA for 3 days (samples are in triplicate). Bar graphs below the blots show band quantitation of Cyp7b1 blot after baseline correction. (D, E & F) Expression of *Abca1* (D), *Ldlr* (E) and *Apoe* (F) mRNA was assessed by qPCR in astrocytes transfected with control siRNA (black bars) or Cyp7b1 siRNA (grey bars) followed by treatment with 0, 1μM or 2μM 25HC for 2 days. Data is normalized to *Actb* (actin B) as the endogenous control gene and shown as relative expression compared to control siRNA with no 25HC. (*0.05-0.01; **0.01-0.001; ***<0.001; ****<0.0001; Two-way ANOVA with Sidak’s multiple comparisons test). (G) ApoE in the conditioned media of astrocytes transfected with control siRNA (black bars) or Cyp7b1 siRNA (grey bars) followed by treatment with 0, 1μM or 2μM 25HC for 2 days. (*0.05-0.01; **0.01-0.001; ***<0.001; ****<0.0001; Two-way ANOVA with Sidak’s multiple comparisons test)

Next, we examined how the knockdown of *Cyp7b1* influences the effect of 25HC. Since the effect of 25HC on the expression of *Abca1* and *Ldlr* waned by 4 days of treatment (Fig 5D & 5E) presumably due to its conversion to 7α25diHC, we tested whether knockdown of *Cyp7b1* can restore the effect of 25HC at 4 days of treatment. In control siRNA cells, the effect of 25HC on *Abca1* expression was minimal by 4 days but was restored by the reduction in Cyp7b1 activity (Fig 6D). Similarly, 25HC was more effective in reducing *Ldlr* expression at 4 days in *Cyp7b1* knockdown astrocytes than in controls (Fig 6E). Expression of *Apoe* was unaffected with or without 25HC in control or *Cyp7b1* siRNA cells (Fig 6F). Despite a general reduction in extracellular ApoE (presumably due to the lipid-based transfection reagent used to transfect the siRNA), astrocytes treated with *Cyp7b1* siRNA secreted 30-40% more ApoE as a result of 25HC treatment (Fig 6G). These results suggest that Cyp7b1 functions as an endogenous regulator of 25HC levels in astrocytes to limit its effects on astrocyte lipid metabolism.

### 25HC-mediated upregulation of LXR-mediated and suppression of SREBP-mediated gene expression are important for ApoE secretion

To determine whether the effects of 25HC on LXR- and SREBP-mediated gene expression are responsible for the increase in extracellular ApoE, we tested the effect of drugs that interfere with each pathway (Fig 7A). To inhibit the effect of 25HC on the LXR pathway, we treated astrocytes with 25HC in the presence of the LXR antagonist, GSK2033 (35). In the presence of GSK2033, very little expression of *Abca1* was observed with or without 25HC treatment (Fig 7B) supporting earlier observations that *Abca1* expressed in astrocytes is LXR-dependent. Interestingly, LY295427 reversed 25HC-dependent upregulation of *Abca1* expression (Fig 7B). Expression of *Ldlr* was suppressed by 25HC as expected and this suppression was unaffected by the presence of GSK2033 (Fig 7C). To interfere with 25HC’s ability to suppress the SREBP pathway, we tested LY295427, a hypocholesterolemic agent that reduces plasma cholesterol levels by increasing *Ldlr* expression in the liver (36, 37). LY295427 also reversed 25HC-dependent suppression of *Ldlr* expression (Fig 7C). These effects of LY295427 may be related to increased expression of Insig1 as previously observed in cells treated with LY295427 (36). Neither GSK2033 nor LY295427 had any effect on *Apoe* gene expression (Fig 7D). However, when the level of ApoE measured in the conditioned media was monitored by ELISA (Fig 7E), both drugs reduced ApoE levels to about 50% of that of control (vehicle-treated) astrocytes. On the other hand, in the presence of 25HC, both drugs eliminated 25HC-stimulated ApoE levels. These results suggest that interfering with either the LXR pathway or the SREBP pathway eliminates the stimulatory effect of 25HC on extracellular ApoE levels in astrocytes.

**Figure 7:**
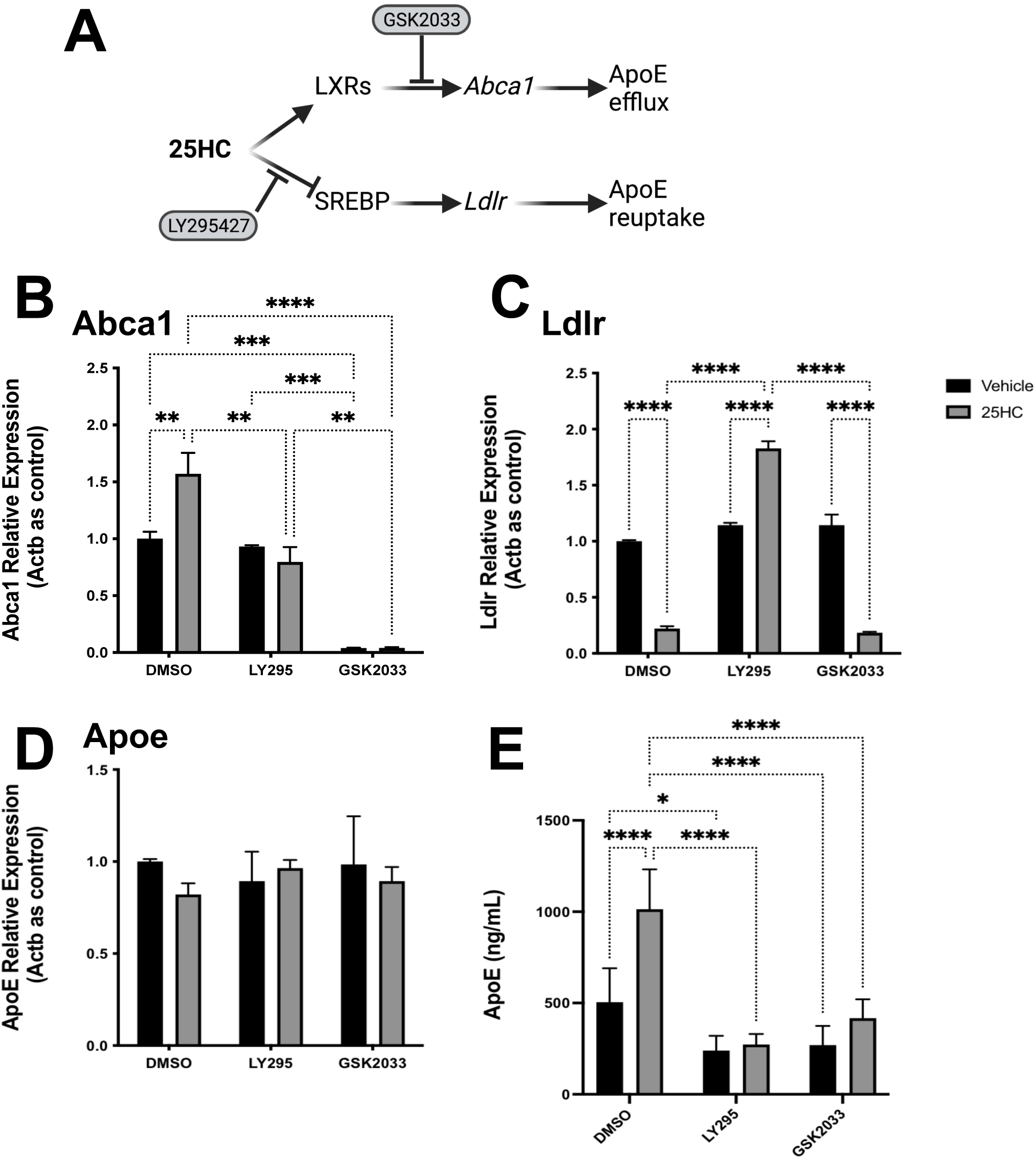
LXR and SREBP pathways are important for increased extracellular ApoE in 25HC-treated astrocytes. (A) Schematic diagram showing the action of inhibitors to block the effect of 25HC on LXR and SREBP pathways. (B, C & D) Expression of *Abca1* (B), *Ldlr* (C) and *Apoe* (D) mRNA in response to treatment of astrocytes with vehicle or 25HC (2μM) in the presence of vehicle (DMSO), 5μM LY295427 or 5μM GSK2033 was assessed by qPCR. Data is normalized to *Actb* (actin B) as the endogenous control gene and shown as relative expression compared to ‘DMSO’. (*0.05-0.01; **0.01-0.001; ***<0.001; ****<0.0001; Two-way ANOVA with Sidak’s multiple comparisons test). (E) ApoE in the conditioned media of astrocytes treated with vehicle or 25HC (2μM) in the presence of vehicle (DMSO), 5μM LY295427 or 5μM GSK2033 was measured by ELISA in the conditioned media after 2 days. (*0.05-0.01; **0.01-0.001; ***<0.001; ****<0.0001; Two-way ANOVA with Sidak’s multiple comparisons test)

### Effect of 25HC on expression of sterol and fatty acid biosynthesis genes in astrocytes

Astrocytes are a major producer of cholesterol and other lipids in the CNS. 25HC is a well-characterized oxysterol regulator of the SREBP pathway in peripheral tissues. However, its importance in the CNS has not been studied. We examined how 25HC influences lipid and cholesterol biosynthetic genes in astrocytes compared to a synthetic LXR agonist, T0901317 (Fig 8). Expression of *Srebf2* (encoding SREBP2, the transcription factor regulating sterol synthesis genes) was reduced to 50% following 25HC treatment of astrocytes for 1 day, whereas T0901317 did not affect *Srebf2* expression (Fig 8A). Similarly, while 25HC also reduced the expression of *Insig1* (a major regulator of the SREBP pathway) to about 20%, T0901317 marginally increased *Insig1* (Fig 8B). Interestingly, while 25HC had no effect on the expression of *Srebf1* (encoding SREBP1, the transcription factor regulating fatty acid synthesis genes), T0901317 increased *Srebf1* by about 50% (Fig 8C). Likewise, while 25HC did not affect the expression of *Fasn* (fatty acid synthase), T0901317 increased *Fasn* by 250% (Fig 8D). 25HC did not affect the expression of *Srebf1* and *Insig2* at 1 day or 4 days of treatment.

**Figure 8:**
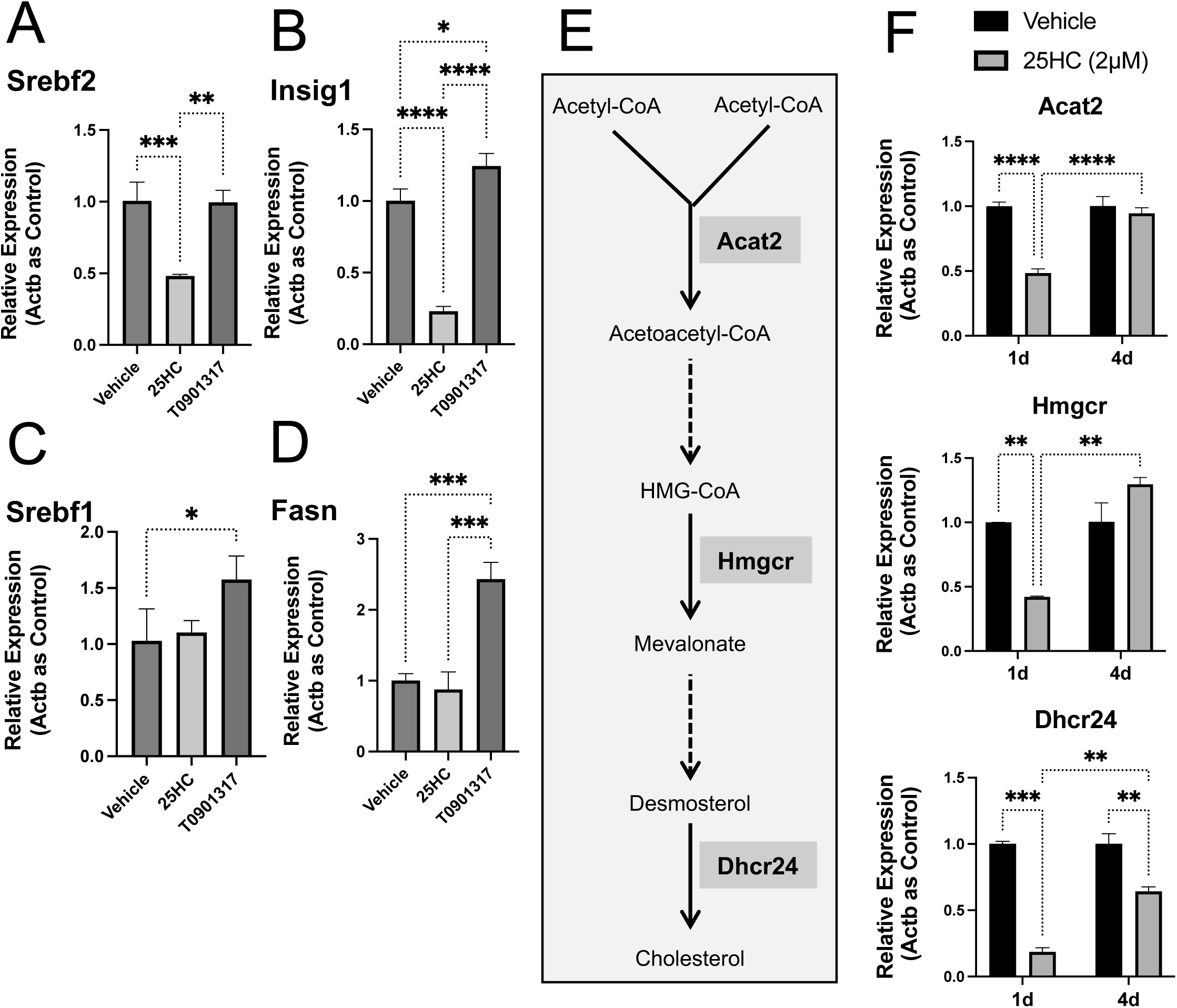
25HC suppresses sterol biosynthesis genes in astrocytes. Expression of *Srebf2* (A), *Insig1* (B), *Srebf1* (C) and *Fasn* (D) mRNA in response to treatment of astrocytes with vehicle, 25HC (2μM) or T0901317 (2μM) for 1 day was assessed by qPCR. Data was normalized to *Actb* (actin B) as the endogenous control gene and shown as relative expression compared to ‘vehicle’. (E) Schematic showing selected steps of cholesterol biosynthesis and the enzyme involved. (F) Expression of *Acat2*, *Hmgcr*, and *Dhcr24* in response to treatment of astrocytes with vehicle (black bars) or 2µM 25HC (grey bars) for 1 day (1d) or 4 days (4d) was quantified by qPCR. Data was normalized to *Actb* (actin B) as the endogenous control gene and shown as relative expression compared to ‘vehicle’. (*0.05-0.01; **0.01-0.001; ***<0.001; ****<0.0001; Two-way ANOVA with Sidak’s multiple comparisons test).

We next examined the expression of three key genes - namely *Acat2*, *Hmgcr* and *Dhcr24* -coding for enzymes required for cholesterol biosynthesis as outlined in Fig 8E. *Acat2* codes for acetyl-coA acyltransferase-2, which catalyzes the first step of cholesterol synthesis. *Hmgcr* codes for 3-hydroxy-3-methylglutaryl-coA reductase, an enzyme catalyzing a key step in the synthesis of mevalonate. *Dhcr24* codes for 24-dehydrocholesterol reductase, which catalyzes the last step in cholesterol biosynthesis. The sterol synthesis genes *Acat2, Hmgcr* and *Dhcr24* were markedly decreased after 1 day of treatment with 25HC, but the effect was diminished by 4 days.

### Changes in astrocytes cholesterol synthesis and esterification elicited by 25HC treatment

Considering the dramatic changes induced by 25HC in cholesterol biosynthesis genes, we next examined whether the changes in gene expression also resulted in corresponding changes in lipids. Astrocytes treated with 25HC or vehicle for 2 days were tested for changes in free fatty acids, free cholesterol and cholesteryl esters (Fig 9). No difference was observed in free fatty acid levels (Fig 9A) in agreement with a lack of effect of 25HC on *Srebf1* and *Fasn* (Fig 8C &D). On the other hand, free cholesterol levels were reduced by nearly 10% in 25HC-treated astrocytes relative to vehicle-treated astrocytes (Fig 9B) reflecting the strong reductions in the expression of *Srebf2* and cholesterol biosynthetic enzymes (Fig 8E & F). Interestingly, the amount of cholesteryl esters (CE) dramatically increased in astrocytes treated with 25HC relative to vehicle (Fig 9C). Deeper analysis of CE revealed that 25HC significantly increased the amounts of CE14:0, CE16:1, CE16:0, CS18:2, CE18:1, CE20:4 and CE22:5, suggesting that 25HC promoted cholesterol esterification with both saturated as well as mono- or poly-unsaturated fatty acids. Previous studies have shown that 25HC activates cholesterol esterification by sterol-O-acyl transferase (SOAT; also known as acyl-coA cholesterol acyltransferase, ACAT). We tested whether 25HC also upregulated expression of the two SOATs - *Soat1* and *Soat2*. While 25HC did not affect expression levels (relative to vehicle control) of *Soat1* (Fig 9E), it increased expression of *Soat2* after 4 days of treatment (Fig 9F). However, the average CT values for *Soat2* were very high (>35) suggesting the expression levels were very low, if any (Fig 9G). The low *Soat2* expression levels and higher *Soat1* expression levels agree with publicly available databases for expression of these genes in the brain (34).

**Figure 9.**
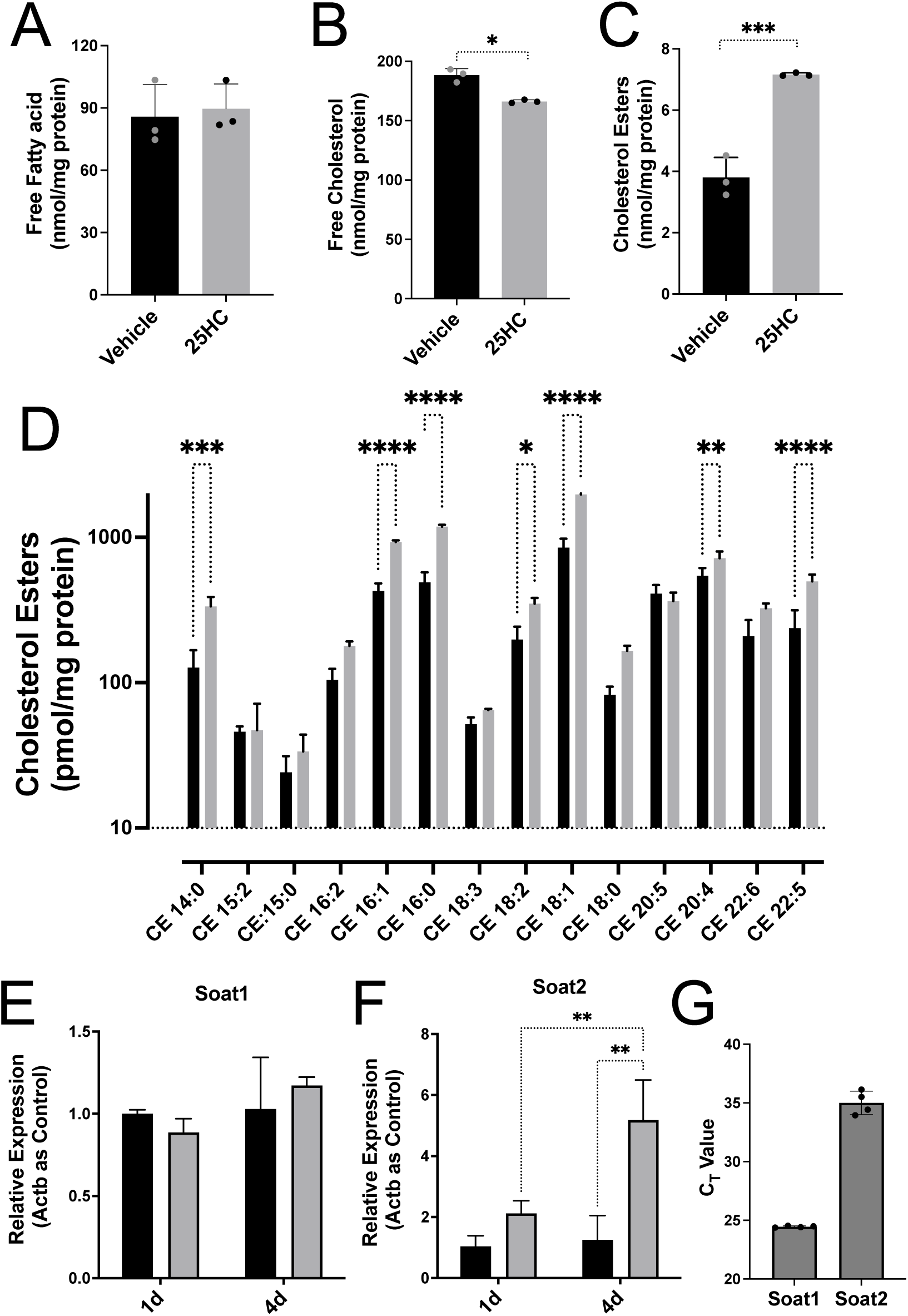
Changes elicited by 25HC in astrocyte lipid profiles. Quantitation of total free fatty acids (A), free cholesterol (B) and total cholesterol esters (C) in mouse astrocytes treated with vehicle (black bars) or with 2µM 25HC (grey bars). Detailed analyses of cholesteryl esters (D) are shown. Expression of *Soat1* (E) and *Soat2* (F) in astrocytes treated with vehicle (black bars) or 2µM 25HC (grey bars) was measured by qPCR and normalized to *Actb* as before. (*0.05-0.01; **0.01-0.001; ***<0.001; ****<0.0001; Data compared with unpaired t-tests in A. For all others two-way ANOVA with Sidak’s multiple comparisons test).

### Augmentation of lipid droplets by 25HC is blocked by an inhibitor of sterol-o-acyltransferase

Since cholesterol esters are mostly stored in lipid droplets, we hypothesized that 25HC might promote lipid droplet (LD) biogenesis in astrocytes. To test this, we treated astrocytes with 25HC in the absence or presence of oleic acid (OA), a well-studied enhancer of LD biogenesis as well as with avasimibe, an inhibitor of SOAT/ACAT (Fig 10A, B & C). LDs were detected using BODIPY 493/503 dye which stains neutral lipids. In the absence of OA, 25HC increased LD counts in astrocytes and this increase was reversed by avasimibe (Fig 10A & B) suggesting that the increased cholesteryl esters in 25HC-treated astrocytes were stored in lipid droplets. OA provides the acyl groups required for the formation of triacylglycerols (TAGs) from diacyl glycerides (DAGs) as well as for the formation of cholesteryl esters, which are key neutral lipid components of LDs. Incubation of astrocytes with OA, increased LDs by nearly 10-fold (compare vehicle control in Fig 10B with OA in Fig 10C). Interestingly, coincubation of astrocytes with OA and 25HC, nearly doubled the LD content from that resulting from OA alone (Fig 10A & C). This suggests that in 25HC-treated astrocytes OA further enhanced cholesteryl esters. While avasimibe had no effect on LDs induced by OA alone, it eliminated the LD-stimulatory effect of 25HC and reverted the levels to that of OA alone. These results suggest that cholesteryl esters produced in response to treatment of astrocytes with 25HC were stored in LDs at basal levels or when co-treated with an acyl-donor such as OA.

**Figure 10.**
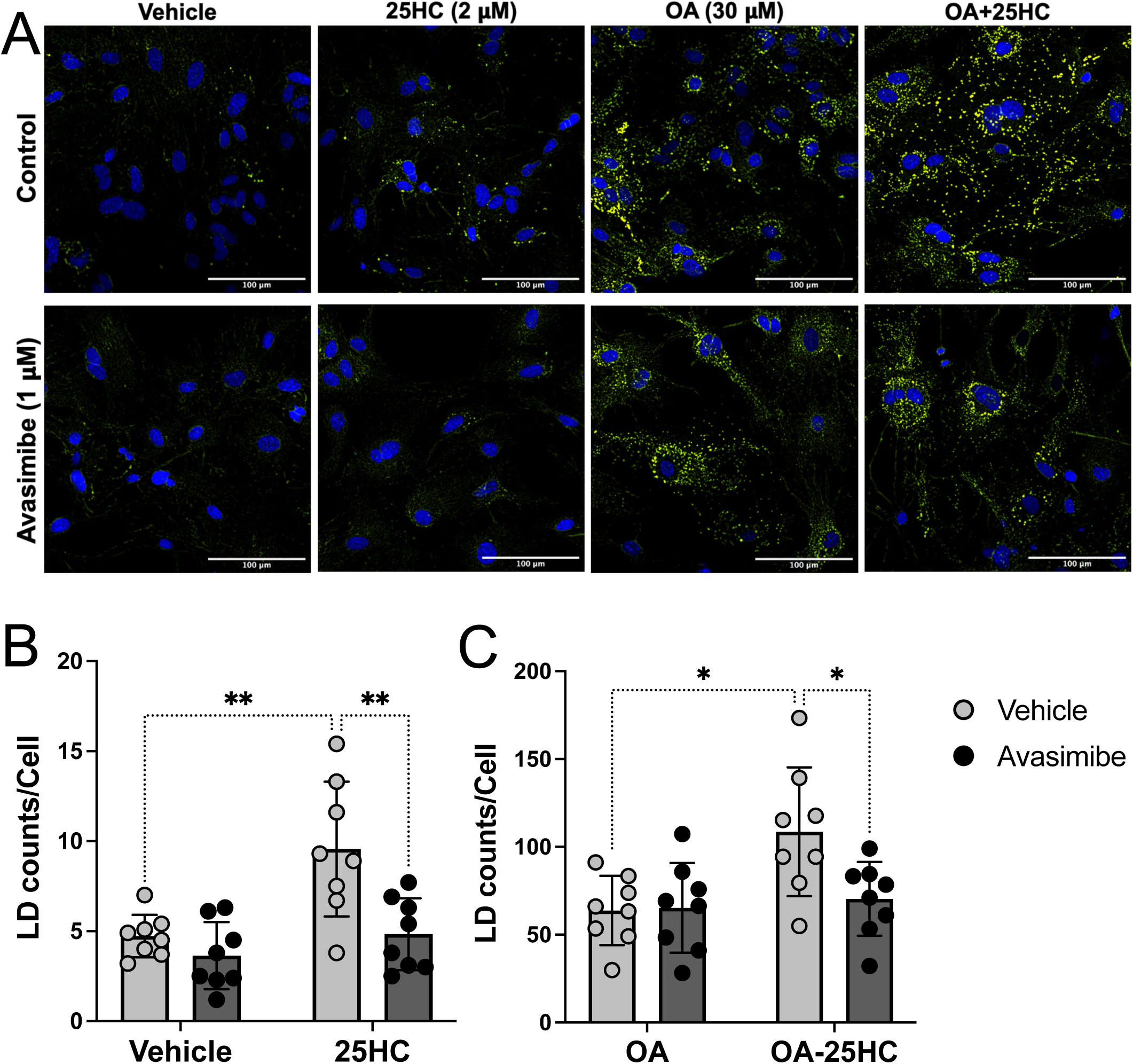
Storage of cholesterol esters in lipid droplets is altered by 25HC. (A) Mouse astrocytes treated with vehicle (ethanol), 2µM 25HC, 30µM OA or OA+25HC were stained for lipid droplets with BODIPY (green) and DAPI (blue). Top row of images is without the SOAT/ACAT inhibitor, avasimibe and bottom row had 1µM avasimibe. Bar = 100µm. (B & C) Quantification of the microscopy data in A and expressed as total LD counts/total nuclei. Light grey bars are without avasimibe and dark grey bars are with avasimibe. (*0.05-0.01; **0.01-0.001; ***<0.001; ****<0.0001; Two-way ANOVA with Sidak’s multiple comparisons test).

## DISCUSSION

In this study, we have provided evidence that 25HC, a cholesterol metabolite produced almost exclusively by activated microglia alters lipid metabolism in astrocytes (Fig 11). We have shown that 25HC alters astrocyte lipid homeostasis in various ways including altering cholesterol metabolism, lipoprotein efflux and reuptake as well as lipid droplet accumulation. Following treatment of mouse astrocytes with physiologically relevant concentrations of 25HC (38), increased secretion as well as decreased reuptake of ApoE lipoproteins resulted in higher levels of extracellular ApoE. Genes required for cholesterol biosynthesis were also strongly downregulated in astrocytes by 25HC. These effects resulted from upregulation of the LXR-mediated and downregulation of SREBP-mediated gene expression by 25HC. 25HC also led to a marked increase in cholesteryl ester levels and its storage in lipid droplets within astrocytes.

**Figure 11:**
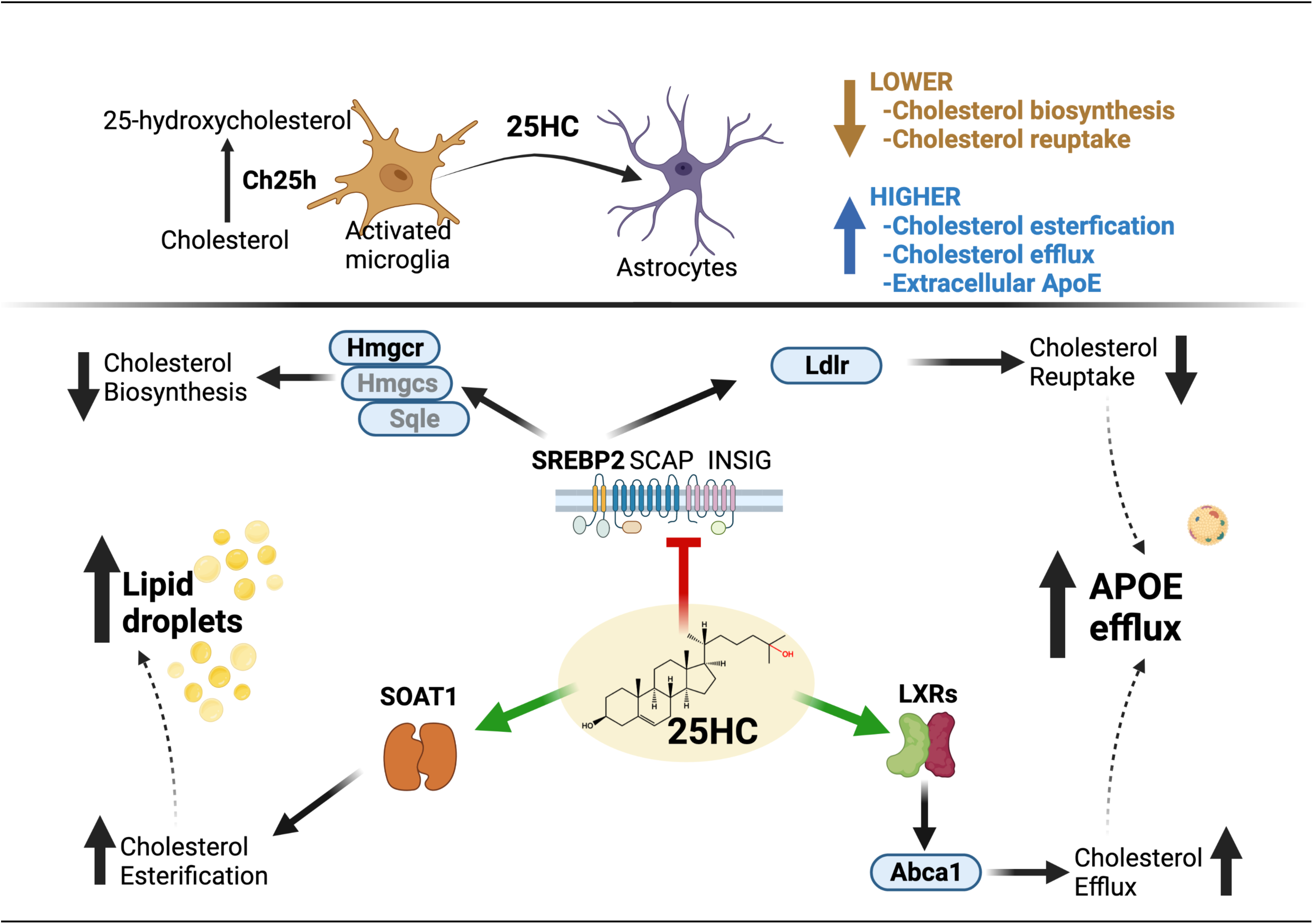
Lipid metabolism and ApoE secretion in astrocytes is modulated by 25HC, a cholesterol metabolite secreted by activated microglia. <u>Top</u>: Overexpression of cholesterol 25-hydroxylase (Ch25h) in microglia activated by inflammatory stimuli converts cholesterol to 25-hydroxycholesterol (25HC). Paracrine effects of secreted 25HC affects astrocyte lipid metabolism and ApoE secretion. <u>Bottom</u>: Schematic showing different ways 25HC alters cholesterol metabolism in astrocytes. 25HC inhibits the SREBP2 pathway to suppress cholesterol biosynthesis by reducing the expression of cholesterol biosynthetic enzymes such as Hmgcr. By suppressing LDL-receptor (Ldlr), 25HC also reduces cholesterol reuptake. Activation of LXRs by 25HC promotes expression of Abca1 that increase cholesterol efflux. Decreased reuptake of lipoproteins as well as increased secretion of lipoproteins result in higher levels of extracellular ApoE. Finally, 25HC enhances the activity of the cholesterol acyl transferase enzyme, Acat1 to promote cholesterol esterification. Thus, 25HC effectively reduces free cholesterol in astrocytes. This figure was made using BioRender.

It is unclear whether 25HC is an endogenous ligand for LXRs *in vivo.* While a recent study showed that 25HC may be an endogenous ligand for LXRs (39), it has been reported to bind LXR weakly (17). However, we have shown that astrocytes readily take up and accumulate 25HC suggesting that the intracellular concentrations of 25HC may be relatively high. Expression of *Apoe* is controlled by LXRs (40) in a human astrocytoma cell line (CCF-STTG1), where synthetic LXR agonists as well as oxysterols promote ApoE gene expression (41, 42). However, our studies (and others) with primary mouse astrocytes show that *Apoe* gene expression is only weakly stimulated by the synthetic LXR agonist, T0901317. Further, *Apoe* expression is neither elevated by 25HC treatment nor inhibited by treatment with the LXR antagonist, GSK2033. This suggests that *Apoe* expression in mouse primary astrocytes does not appear to be strongly dependent on LXRs. However, we observed that 25HC significantly increases the expression of *Abca1*, a well-known LXR target gene (43) and that blocking LXR activity with GSK2033 reduced *Abca1* expression in the presence or absence of 25HC.

Astrocytes are a major cell type in the brain for cholesterol biosynthesis, which is controlled by *Srebf2* (18) and may play an important role in the pathogenesis of AD. Cholesterol biosynthesis controlled by the SREBP-pathway encompasses the following key steps: (i) when cholesterol levels are low Scap-Srebp2 complex migrates from the ER to the Golgi via COP-II-dependent trafficking; (ii) in the Golgi Srebp2 is proteolytically cleaved by the S1P and S2P proteases to generate the cytosolic Srebp2 fragment; (iii) the transcriptionally active Srebp2 fragment translocates to the nucleus to stimulate transcription of cholesterol synthesis genes. Polymorphisms in the *SREBF2* gene (rs2269657) have been reported to be associated with the risk of developing late onset AD and *SREBF2* mRNA was reported to be significantly higher in AD temporal cortex and cerebellum (44). Astrocyte-derived cholesterol was reported to be important for beta-amyloid production in neurons (45). Further, nuclear Srebp2 has been reported to be negatively correlated with the presence of AT8-positive neurofibrillary tangles in AD brain (46). In mice, astrocyte-specific loss of *Srebf2* resulted in reduced cholesterol synthesis and defects in neurite outgrowth, behavior, and energy metabolism (47). When intracellular cholesterol or oxysterol levels are high, cholesterol synthesis is turned down (18). The binding of 25HC to the Insig-Scap interface retains the Scap-Srebp2 complex in the ER precluding its proteolytic processing in the Golgi (48). Treatment of primary astrocytes with 25HC resulted in markedly reduced expression of *Insig1* as well as *Srebf2*, which encodes the Srebp2 protein itself. When 25HC binds to Insig1, it also stabilizes the protein by reducing proteasomal degradation of Insig1 thus promoting its interaction with Scap-Srebp2 (49). Since *Insig1* is a target of the Srebp-signaling pathway (49), expression of *Insig1* is also reduced in the presence of 25HC. This intricate control of *Insig1* gene expression and protein stability (and function) establishes a feedback mechanism by which cholesterol biosynthesis is regulated by 25HC.

Following treatment of mouse astrocytes with 25HC, we observed a dramatic decrease in the expression of *Ldlr*, a gene expressed under the control of the SREBP pathway. Ldlr is a major receptor for the reuptake of ApoE lipoproteins (50). Ldlr is normally localized to the plasma membrane where it binds extracellular ApoE lipoproteins followed by endocytosis and subsequent release of bound lipids in the lysosome (51). Overexpression of LDLR reduced ApoE levels and reduced tau-mediated neurodegeneration (52). On the other hand, by reducing Ldlr levels, 25HC markedly limits reuptake of ApoE resulting in greater amounts of extracellular ApoE. A previous study showed that LY295427 reverses the effect of 25HC on the SREBP-pathway by elevating the expression of *Ldlr* (53) as well as *Insig1* (36) presumably by creating a sink for 25HC. In agreement, we observed that 25HC failed to suppress *Ldlr* expression in astrocytes treated with LY295427. However, it is unclear whether LY295427 also reverses the suppression of *Srebf2* expression by 25HC. Taken together, microglial 25HC overproduced under neuroinflammatory conditions may be a potentially important feedback regulator of the SREBP machinery in astrocytes.

25HC also nearly doubled levels of cholesterol esters and their storage in LDs in astrocytes. The increase in 25HC-induced LDs could be blocked by an inhibitor of SOAT/ACAT suggesting the accumulation in CE-rich lipid droplets in 25HC-treated astrocytes. Unlike triglyceride-enriched LDs, CE-enriched LDs are though to play an important role in the biosynthesis of steroid hormones (54). It will be interesting to examine whether mice lacking *Ch25h* are defective in the production of neurosteroids.

With increased cholesterol efflux, increased cholesterol esterification and decreased cholesterol biosynthesis, the net effect of 25HC on astrocytes is a reduction in overall free cholesterol levels by about 10% in astrocytes as we empirically established. Over long periods, these cholesterol-reducing effects are likely to be detrimental to cell survival. However, our results show that the effects of 25HC are lower when treated over 4 days compared to treatments for 1 day. 25HC is metabolized to 7α,25-dihydroxycholesterol (7α25diHC) by the P450 enzyme, Cyp7b1 (55) and is subsequently converted to bile acid for removal (56). Interestingly, 7α25diHC does not appear to interfere with astrocyte lipid metabolism as there was no effect of 7α25diHC on extracellular ApoE levels in our astrocyte cultures.

However, 7α25diHC is a chemoattractant ligand for Gpr183/Ebi2 expressed on lymphocytes (57). It has been previously shown that brain levels of 25HC are dramatically increased in *cyp7b1*-/-mice (58). Loss of function mutations in *CYP7B1* cause autosomal recessive spastic paraplegia 5A (59), a motor neuron disease wherein defective cholesterol homeostasis is implicated (60). Our data show that astrocytic Cyp7b1 serves to limit the effects of 25HC and when *Cyp7b1* mRNA was knocked down using RNAi, the effects of 25HC on increased *Abca1* expression and reduced *Ldlr* expression were prolonged as was the increase in extracellular ApoE.

The results presented herein highlight a potential role for 25HC, the immune-oxysterol produced by microglia to regulate extracellular levels of ApoE as well as cholesterol metabolism in astrocytes. We postulate that 25HC may function as a potential mediator of microglia-astrocyte crosstalk and a potent regulator of astrocyte lipid metabolism under conditions of neuroinflammation and accompanying microglial activation. It will be interesting to examine the effects 25HC in in vivo models of neuroinflammation.

### DATA AVAILABILITY

All data are contained within the manuscript.

## Abbreviations

25HC: 25-hydroxycholesterol
Ch25h: cholesterol 25-hydroxylase
AD: Alzheimer disease
7α25diHC: 7α, 25-dihydroxycholesterol
LPS: lipopolysaccharide
qPCR: quantitative polymerase chain reaction
BODIPY: boron-dipyrromethene
FC: Free cholesterol
CE: Cholesteryl ester
LD: lipid droplets
SOAT: sterol-O-acyltransferase

## SUPPORTING INFORMATION

This article contains supporting information.

## ACKNOWLEDGEMENTS

We wish to acknowledge the gift of LY295427 from Dr. Doug Covey, Department of Developmental Biology, Washington University School of Medicine, St Louis, MO.

## AUTHOR CONTRIBUTIONS

AGC and SMP, conceived the study; AGC, SMP and DMH, obtained funding; AGC, DTR, DT, JR and XH, performed experiments and analyzed data; DTR, JML, DMH and SMP, critically reviewed the manuscript; AGC, wrote the manuscript

---

**Supplementary Figure 1:**
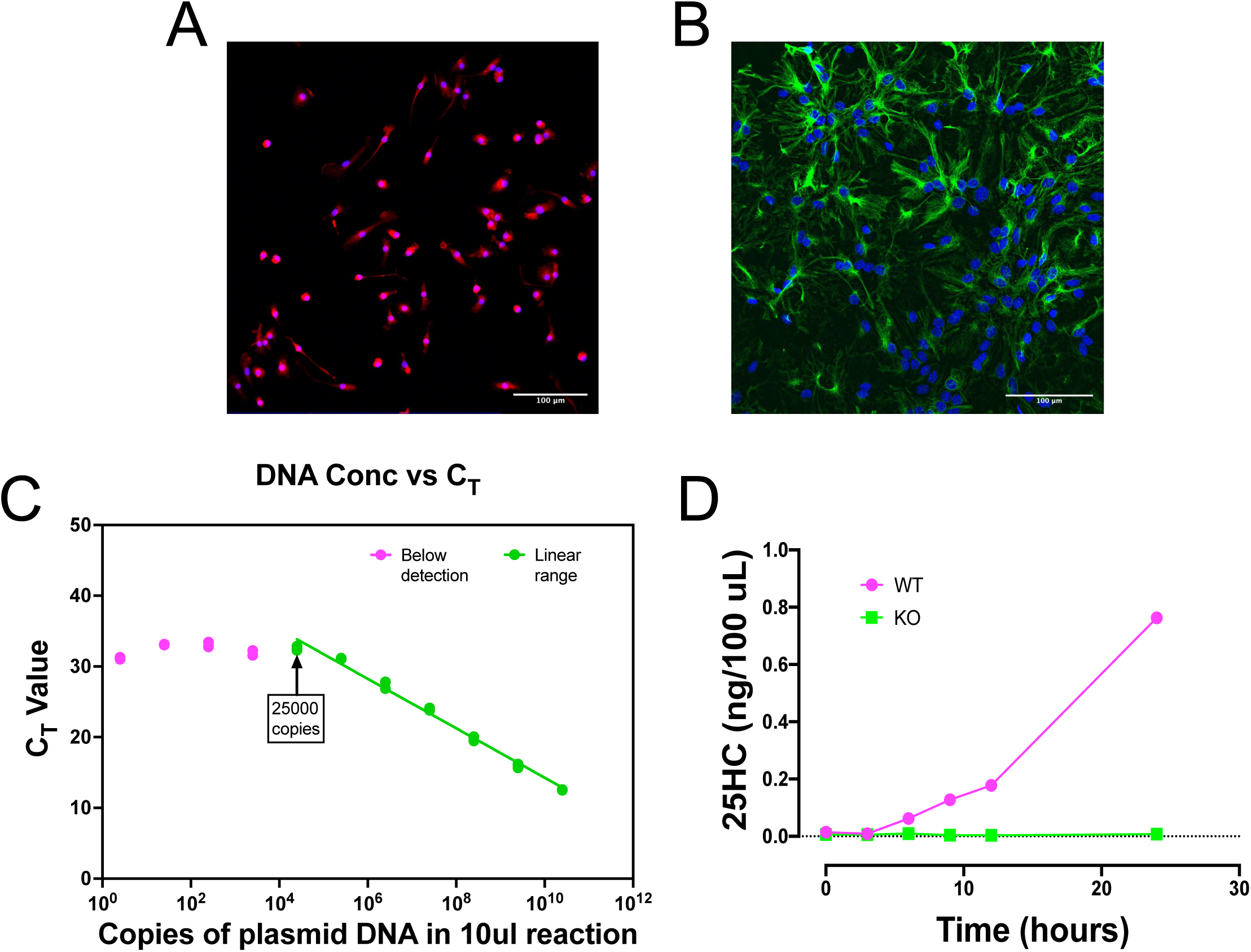
Supplementary data corresponding to Figure 1. Purity of representative cultures of microglia (A) and astrocytes (B) used in this study. Microglial cultures were stained for Iba1 (red) and DAPI (blue). Astrocyte cultures were stained for GFAP (green) and DAPI (blue). Bar indicates 100µm. (C) Plot of the standard curve for CT values versus DNA concentration. A purified plasmid carrying the mouse *Ch25h* gene was in 10-fold serial dilutions and a qPCR was performed to generate this plot. CT values decreased linearly when there were 25000 or more copies the *Ch25h* gene in a 10μl reaction. CT values below 25000 copies were non-linear. CT values obtained for cDNA prepared from 200,000 microglia or astrocytes were in the linear range of the plot. (D) Ch25h-dependent production and secretion of 25HC by microglia was affirmed by testing 25HC in the conditioned media of wildtype and ch25h -/- microglia treated with LPS for various times. No 25HC was detectable in ch25h-/- microglia.

